# Reward learning over weeks versus minutes increases the neural representation of value in the human brain

**DOI:** 10.1101/158964

**Authors:** G. Elliott Wimmer, Jamie K. Li, Krzysztof J. Gorgolewski, Russell A. Poldrack

## Abstract

Over the past few decades, neuroscience research has illuminated the neural mechanisms supporting learning from reward feedback. Learning paradigms are increasingly being extended to study mood and psychiatric disorders as well as addiction. However, one potentially critical characteristic that this research ignores is the effect of time on learning: human feedback learning paradigms are usually conducted in a single rapidly paced session, while learning experiences in ecologically relevant circumstances and in animal research are almost always separated by longer periods of time. In our experiments, we examined reward learning in short condensed sessions distributed across weeks vs. learning completed in a single “massed” session in male and female participants. As expected, we found that after equal amounts of training, accuracy was matched between the spaced and massed conditions. However, in a 3-week follow-up, we found that participants exhibited significantly greater memory for the value of spaced-trained stimuli. Supporting a role for short-term memory in massed learning, we found a significant positive correlation between initial learning and working memory capacity. Neurally, we found that patterns of activity in the medial temporal lobe and prefrontal cortex showed stronger discrimination of spaced-vs. massed-trained reward values. Further, patterns in the striatum discriminated between spaced-and massed-trained stimuli overall. Our results indicate that single-session learning tasks engage partially distinct learning mechanisms from spaced sessions of training. Our studies begin to address a large gap in our knowledge of human learning from reinforcement, with potential implications for our understanding of mood disorders and addiction.

**Significance statement:** Humans and animals learn to associate predictive value with stimuli and actions, and these values then guide future behavior. Such reinforcement-based learning often happens over long time periods, in contrast to most studies of reward-based learning in humans. In experiments that tested the effect of spacing on learning, we found that associations learned in a single massed session were correlated with short-term memory and significantly decayed over time, while associations learned in short massed sessions over weeks were well-maintained. Additionally, patterns of activity in the medial temporal lobe and prefrontal cortex discriminated the values of stimuli learned over weeks but not minutes. These results highlight the importance of studying learning over time, with potential applications to drug addiction and psychiatry.

## Introduction

When making a choice between an apple and a banana, our decision often relies on values shaped by countless previous experiences. By learning from the outcomes of these repeated experiences, we can make efficient and adaptive choices in the future. Over the past few decades, neuroscience research has revealed the neural mechanisms supporting this kind of learning from reward feedback, demonstrating a critical role for the striatum and the midbrain dopamine system (Houk et al., 1995; Schultz et al., 1997; Dolan and Dayan, 2013; Steinberg et al., 2013). However, research in humans has tended to focus on two different timescales: short-term learning from reward feedback across minutes, for example, in “bandit” or probabilistic selection tasks (Frank et al., 2004; Daw et al., 2006), or choices based on well-learned values, for example, over snack foods (Plassmann et al., 2007). There has been remarkably little research in humans that examines how value associations are maintained or acquired beyond a single session (Herbener, 2009; Tricomi et al., 2009; Grogan et al., 2017; de Wit et al., in press), even though our preferences are often shaped across multiple days, months, or years of experience.

Recently, researchers have begun to use learning tasks in combination with reinforcement learning models to investigate behavioral dysfunctions in mood and psychiatric disorders as well as addiction in the growing area of “computational psychiatry” (Maia and Frank, 2011; Schultz, 2011; Montague et al., 2012; Whitton et al., 2015; Moutoussis et al., 2016). This translational work on human reward-based learning builds on research in animals where circuit functions can be extensively manipulated (Steinberg et al., 2013; Ferenczi et al., 2016). However, at the condensed timescale of most human paradigms, “massed” timing likely allows processes in addition to dopaminergic mechanisms of feedback-based learning, such as working memory, to support behavior (Collins and Frank, 2012; Collins et al., 2014).

While no studies have directly compared values learned in a massed session or across days, several recent neuroimaging studies have examined the neural representation of values learned across days (Tricomi et al., 2009; Wunderlich et al., 2012), supporting a role for the human posterior striatum in representing the value of well-learned stimuli. These findings align with neurophysiological recordings in the striatum of animals (Yin and Knowlton, 2006; Kim and Hikosaka, 2013). However, reward-related BOLD responses in the putamen are not selective to consolidated or “habitual” reward associations (e.g. O’Doherty et al., 2003; Dickerson et al., 2011; Wimmer et al., 2014); moreover, previous studies did not allow for a matched comparison between newly-learned reward associations and consolidated associations.

In addition to the striatum, fMRI and neurophysiological studies have shown that responses in the medial temporal lobe and hippocampus are correlated with reward and value (Lebreton et al., 2009; Wirth et al., 2009; Lee et al., 2012; Wimmer et al., 2012). While these responses are not easily explained by a relational memory theory of MTL function (Eichenbaum and Cohen, 2001), they may fit within a more general view of the hippocampus in supporting some forms of statistical learning (including stimulus-stimulus associations; Schapiro et al., 2012; Schapiro et al., 2014). Memory mechanisms in the MTL may also play a role in representing previous episodes that can be sampled to make a reward-based decision (Murty et al., 2016; Wimmer and Buechel, 2016; Bornstein et al., 2017), a role that could be enhanced by consolidation.

To characterize the cognitive and neural mechanisms which support learning long-term reward associations, we utilized a simple reward-based learning task. Participants initially learned value associations for spaced stimuli in the lab and then online across two weeks in multiple short massed sessions. Associations for massed stimuli were acquired during a second in-lab session over approximately 20 minutes (followed by fMRI scanning in one group), similar to the kind of training commonly used in reinforcement learning tasks. Finally, to examine maintenance of learning, a long-term test was administered three weeks after the completion of training.

## Methods

### Participants and Overview

Participants were recruited via advertising on the Stanford Department of Psychology paid participant pool web portal (https://stanfordpsychpaid.sona-systems.com). Informed consent was obtained in a manner approved by the Stanford University Institutional Review Board. In study 1, behavioral and fMRI data acquisition proceeded until fMRI seed grant funding expired, leading to a total of 33 scanned participants in the reward learning task. In order to ensure that the fMRI sessions two weeks after the first in-lab session were fully subscribed, a total of 62 participants completed the first behavioral session. Of this group, a total of 29 participants did not complete the fMRI and behavioral experiment described below. The results of 33 participants (20 female) are included in the analyses and results, with a mean age of 22.8 years (range: 18-34). Participants were paid $10/hour for the first in-lab session and $30/hour for the second in-lab (fMRI) session, plus monetary rewards from the learning phase and choice test phase.

In Study 2, a total of 35 participants participated in the first session of the experiment, but 4 were excluded from the final dataset, as described below. Our sample size was designed to approximately match the size of Study 1. The final dataset included 31 participants (24 female), with a mean age of 23.3 years (range: 18-32). Two participants failed to complete the second in-lab session and all data were excluded; one other participant exhibited poor performance the first session (less than 54% correct during learning and less than 40% correct in the choice test) and was therefore excluded from participation in the follow-up sessions. Of the 31 included participants, one participant failed to complete the third in-lab session, but data from other sessions were included. Participants were paid $10/hour for the two in-lab sessions, monetary rewards from the learning phase and choice test phase, plus a bonus of $12 for the 5-minute duration third in-lab session.

Both Study 1 and Study 2 utilized the same reward-based learning task (adapted from Gerraty et al., 2014). Participants learned the best response for individual stimuli in order to maximize their payoff. Two different sets of stimuli were either trained in multiple massed sessions spaced across two weeks (“spaced-trained” stimuli) or in a single session (“massed-trained” stimuli; **Fig. 1A**). Initial spaced learning began in the first in-lab session and continued across three online training sessions spread across approximately 2 weeks. Initial learning about massed stimuli began in the second in-lab session and continued until training was complete. Spaced training always preceded massed training, so that by the end of the second in-lab session both sets of stimuli had been shown on an equal number of learning trials. This design was the same across Study 1 and Study 2, with the difference that Study 1 included an fMRI portion at the time of the second in-lab session and that post-learning tests were conducted at different points during learning in the two studies. Additionally, the three-week follow-up measurement was conducted online for Study 1 and in-lab for Study 2.

**Figure 1.**
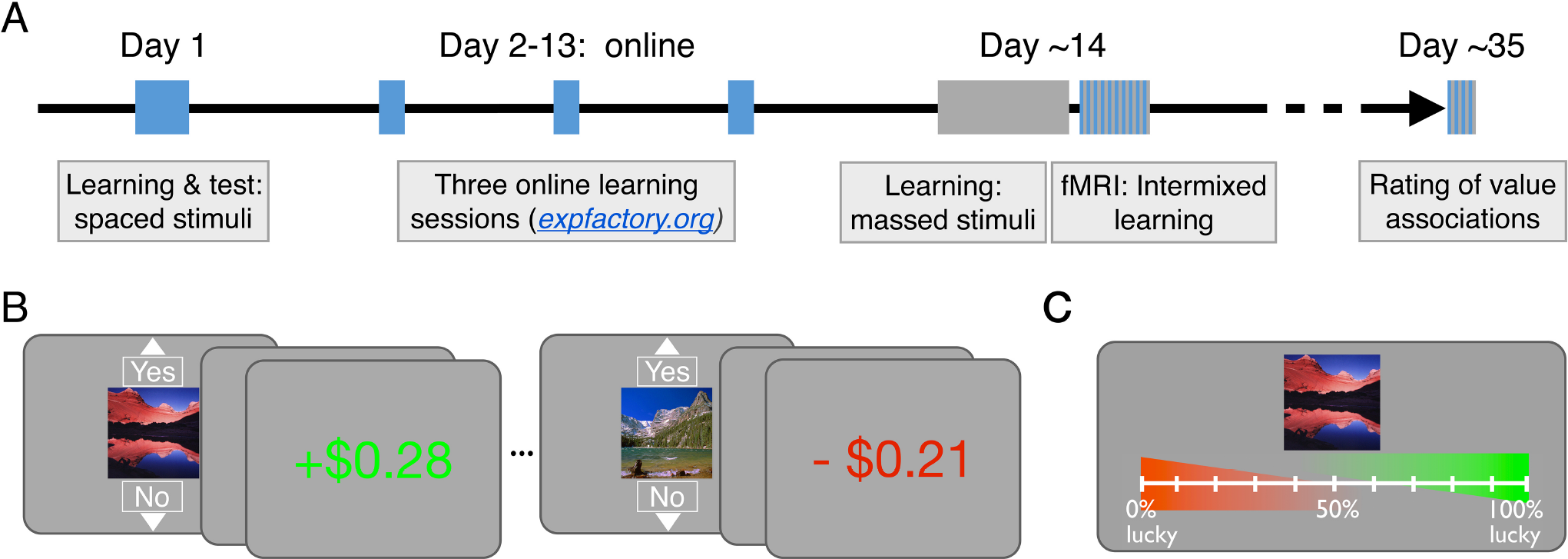
***A***, Experimental timeline. Learning for the spaced-trained stimuli is indicated in blue and learning for the massed-trained stimuli is indicated in grey. The initial learning session for spaced-trained stimuli was completed on day 1. Learning for spaced stimuli was then competed in multiple short (massed) sessions, while learning for massed stimuli was completed in a single session approximately 14 days later. Aside from the separation of spaced learning into multiple condensed sessions, inter-trial timing was matched across conditions. A forced-choice test was also collected after initial learning and the completion of learning. A long-term follow-up measure of reward value using ratings was collected after approximately 3 weeks. ***B***, Reward learning task. Participants learned to select “Yes” for reward-associated stimuli and select “No” for loss-associated stimuli. Choices were presented for 2 sec, and feedback followed after a 1 sec delay. ***C***, Reward association rating test. This rating scale followed the initial in-lab learning sessions and was also administered 3 weeks after the last learning session.

We chose to use a simple instrumental learning task (adapted from Gerraty et al., 2014) instead of a choice-based learning task for three reasons. First, the majority of the animal work we are translating involves relatively simple instrumental or Pavlovian value learning designs with a single focal stimulus (e.g. Schultz et al., 1997; O’Doherty et al., 2003; Tricomi et al., 2009), including previous work on spacing effects in feedback learning (Spence and Norris, 1950; Teichner, 1952). Most directly, the present design was inspired by the work of Hikosaka and colleagues on long-term memory for value (Kim and Hikosaka, 2013; Kim et al., 2015; Ghazizadeh et al., 2018) and by the work of Collins and colleagues on short-term memory contributions to rapid feedback-based learning (Collins and Frank, 2012; Collins et al., 2014). Second, a single-stimulus design avoids differential attentional allocation toward the relatively more valuable stimulus in a set, which is inherent in multi-stimulus designs (Daw et al., 2006; Pessiglione et al., 2006). Third, our reward learning paradigm has been shown previously to effectively establish stimulus-value associations (Gerraty et al., 2014), as evidenced by the ability of newly learned stimulus-reward associations to transfer or generalize across previously established relational associations. Such transfer is related to striatal correlates of learned value (Wimmer and Shohamy, 2012). While our in-task learning measures are related to (“Yes” / “No”) action value, this previous work and recent human fMRI research indicate that mechanisms supporting the learning of stimulus-action and stimulus-value associations operate at the same time (Colas et al., 2017).

### Experimental design, Study 1

In Study 1, before the learning phase, participants rated a set of 38 landscape picture stimuli based on liking, using a computer mouse, preceded by one practice trial. The same selection procedure and landscape stimuli were used previously (Wimmer and Shohamy, 2012). These ratings were used to select the 16 most neutrally-rated set of stimuli per participant to be used in Study 1. Stimuli were then randomly assigned to condition (spaced or massed) and value (reward or loss). In Study 2, we used the ratings collected across participants in Study 1 to find the most neutrally-rated stimuli on average and then created two counterbalanced lists of stimuli from this set.

Next, in the reward game in both studies, participants’ goal was to learn the best response (arbitrarily labeled “Yes” and “No”) for each stimulus. Participants used up and down arrow keys to make “Yes” and “No” responses, respectively. Reward-associated stimuli led to a win of $0.35 on average when “Yes” was selected and a small loss of - $0.05 when “No” was selected. Loss-associated stimuli led to a neutral outcome of $0.00 when “No” was selected and −$0.25 when “Yes” was selected. These associations were probabilistic, such that the best response led to the best outcome 80% of the time during training. If no response was recorded, at feedback a warning was given: “Too late or wrong key! - $0.50”, and participants lost $0.50.

In a single reward learning trial, a stimulus was first presented with the options “Yes” and “No” above and below the image, respectively (**Fig. 1B**). Participants had 2 seconds to make a choice. After the full 2 s choice period, a 1 s blank screen ITI preceded feedback presentation. Feedback was presented in text for 1.5 s, leading to a total trial duration of 4.5 s. Reward feedback above +$0.10 was presented in green, and feedback below $0.00 was presented in red, while other values were presented in white. After the feedback, an ITI of duration 2 preceded the next trial (min, 0.50 s; max, 3.5 s), where in the last 0.25 s prior to the next trial the fixation cross turned from white to black. The background for all parts of the experiment was grey (RGB value [111 111 111]). We specifically designed the timing of feedback (3 sec from the onset of choice) to fall within the range of previous studies on feedback-based learning and the dopamine system which show a strong decay of the fidelity of the dopamine reward prediction error response as feedback is delayed beyond several seconds (Fiorillo et al., 2008); beyond this point, other (e.g. hippocampal) mechanisms may support learning (Foerde et al., 2013).

To increase engagement and attention to the feedback, we introduced uncertainty into the feedback amounts in two ways: first, all feedback amounts were jittered ± $0.05 around the mean using a flat distribution. Second, for the reward-associated stimuli, half were associated with a low reward amount ($0.45) and half with a higher reward amount ($0.25). We did not find that this second manipulation significantly affected learning performance at the end of the training phase, and thus our analyses and results collapse across the reward levels.

In the initial spaced learning session in-lab, participants learned associations for spaced-trained stimuli, which differed from the training for massed-trained stimuli only in that training for spaced stimuli was spread across 4 “massed” sessions, 1 in-lab and 3 online. Initial learning followed by completion of learning for massed-trained stimuli occurred in the subsequent second in-lab session. The spaced-and massed-trained conditions each included 8 different stimuli, of which half were associated with reward and half were associated with loss. In the initial learning session for both conditions, each stimulus was repeated 10 times. The lists for the initial learning session were pseudo-randomized, with constraints introduced to facilitate initial learning and to achieve ceiling performance before the end of training.

In order to more closely match the delay between repetitions commonly found in human studies of feedback-based learning (e.g. Pessiglione et al., 2006), where only several different trial types are included, we staged the introduction of the 8 stimuli into two sets. In the initial learning session for both spaced-and massed-trained stimuli, 4 stimuli were introduced in the first 40 trials and the other 4 stimuli were introduced in the second 40 trials. Further, when a new stimulus was introduced, the first repetition followed immediately. The phase began with 4 practice trials including 1 reward-associated practice stimulus and 1 loss-associated practice stimulus, followed by a question about task understanding. Three rest breaks were distributed throughout the rest of the phase.

After the initial learning session in both conditions, participants completed a reward rating phase and an incentive-compatible choice phase. In the reward rating phase, participants tried to remember whether a stimulus was associated with reward or not. They were instructed to use a rating scale to indicate their memory and their confidence in their memory using a graded scale, with responses made via computer mouse (**Fig. 1C**). Responses were self-paced. After 0.5 s, trials were followed by a 3 s ITI. For analyses, responses (recorded in pixel left-right location values) were transformed to 0-100 percent.

In the incentive-compatible choice phase, participants made a forced-choice response between two stimuli, only including spaced stimuli in the first in-lab session and only including massed stimuli in the second in-lab session. Stimuli were randomly presented on the left and right side of the screen. Participants made their choice using the 1-4 number keys in the top row of the keyboard, with a ‘1’ or ‘4’ response indicating a confident choice of the left or right option, respectively, and a ‘2’ or ‘3’ response indicating a guess choice of the left or right option, respectively. The trial terminated 0.25 s after a response was recorded, followed by a 2.5 s ITI. Responses were self-paced. Participants were informed that they would not receive feedback after each choice but that the computer would keep track of the number of correct choices of the reward-associated stimuli that were made and pay a bonus based on their performance. As the long-term follow-up only included ratings, choice analyses were limited to comparing how choices aligned with ratings.

At the end of the session, participants completed two additional measures. We collected the Beck Depression Inventory (BDI), but scores were too low and lacked enough variability to enable later analysis (median score = 2 out of 69 possible; scores above 13 indicate mild depression). The second measure we collected was the operation-span task (OSPAN), which was used as an index of working memory capacity (Lewandowsky et al., 2010; Otto et al., 2013). In the OSPAN, participants made accuracy judgments about simple arithmetic equations (e.g. ‘2 + 2 = 5’). After a response, an unrelated letter appeared (e.g. ‘B’), followed by the next equation. After arithmetic-letter sequences ranging in length from 4 to 8, participants were asked to type in the letters that they had seen in order, with no time limit. Each sequence length was repeated 3 times. In order to ensure that participants were fully practiced in the task before it began, the task was described in-depth in instruction slides, followed by 5 practice trials. Scores were calculated by summing the number of letters in fully correct letter responses across all 15 trials (mean, 49.9; range, 19-83) (Otto et al., 2013); mean performance on the arithmetic component was 81.9%.

### Online training

Subsequent to the first in-lab session where training on spaced stimuli began, participants completed three short online “massed” sessions with the spaced-trained stimuli. Sessions were completed on a laptop or desktop computer (but not on mobile devices), using the <u>expfactory.org</u> platform (Sochat et al., 2016). Code for the online reward learning phase can be found at: https://github.com/gewimmer-neuro/reward_learning_js. Each online training session included 5 repetitions of the 8 spaced-trained stimuli, in a random order, leading to 15 additional repetitions per spaced-trained stimulus overall. The task and timing were the same as in the in-lab sessions, with the exception that the screen background was white and the white feedback text was replaced with grey. Participants completed the online sessions across approximately 2 weeks, initiated with an email from the experimenter including login details for that session. In the case that participants had not yet completed the preceding online session when the notification about the next session was received, participants were instructed to complete the preceding session that day and the next session the following day. Thus, at least one overnight period was required between sessions. Participants were instructed to complete the session when they were alert and not distracted. We found that data for two sessions in one participant were missing and for an additional 7 participants, data for one online session was missing. Based on follow-up with a subset of participants, we can conclude that missing data was due in some cases to technical failures and in some cases due to non-compliance. Among participants with a missing online spaced training session, performance during scanning for spaced-trained stimuli was above the group mean (94.8% vs. 91.0%). Note that if a subset of participants did not complete some part of the spaced training, this would, if anything, weaken any differences between spaced and massed training.

### Second in-lab session

Next, participants returned for a second in-lab session, approximately two weeks later (mean, 13.5 days; range, 10-20 days). Here, participants began and completed learning on the massed-trained stimuli. Initial training across the first 10 repetitions was conducted as described above for the first in-lab session. Next, participants completed a rating phase including both spaced-and massed-trained stimuli and choice phase involving only the massed-trained stimuli. After this, participants finished training on the massed-trained stimuli, bringing total experience up to 25 repetitions, the same as for the spaced-trained stimuli to that point.

In Study 1, participants next entered the scanner for an intermixed learning session. Across 2 blocks, participants engaged in additional training on the spaced-and massed-trained stimuli, with 6 repetitions per stimulus. With four initial practice trials, there were 100 total trials. During scanning, task event durations were as in the behavioral task above, and ITI durations were on average 3.5 s (min, 1.45 s; max, 6.55 s). Responses were made using a button cylinder, with the response box positioned to allow finger responses to mirror those made on the up and down arrow keys on the keyboard.

Following the intermixed learning session, participants engaged in single no-feedback block, where stimuli were presented with no response requirements. This block provided measures of response to stimuli without the presence of feedback, and lists were designed to allow for tests of potential cross-stimulus repetition-suppression (Barron et al., 2013; Klein-Flugge et al., 2013; Barron et al., 2016). Stimuli were presented for 1.5 s, followed by a 1.25 s ITI (range, 0.3 – 3.7 s). To provide a measure of attention and to promote recollection and processing of stimulus value, participants were instructed to remember whether a stimulus had been associated with reward or with no reward. On ~10% of trials, 1 s after the stimulus had disappeared, participants were asked to answer whether the best response to the stimulus was a “Yes” or a “No”. Participants had a 2 s window in which to make their response; no feedback was provided unless a response was not recorded, in which case the warning “Too late or wrong key! −$0.50” was displayed. Each stimulus was repeated 10 times during the no-feedback phase, yielding 160 trials. Different stimuli of the same type (spaced training by reward value) were repeated on sequential trials to allow for repetition suppression analyses. At least 18 sequential events for each of these critical 4 comparisons were presented in a pseudorandom order.

In Study 1, participants also engaged in an additional unrelated cognitive task during the scanning session (approximately 30 min) and a resting scan (8 min). The order of the cognitive task and the reward learning task were counterbalanced across participants. Results from the cognitive task will be reported separately.

After scanning, participants engaged in an exploratory block to study whether and how participants would reverse their behavior given a shift in feedback contingencies. Importantly, the “reversed” stimuli and control non-reversed stimuli (4 per condition per participant) were not included in the analyses of the 3-week follow-up data. One medium-reward stimulus and one loss-associated stimulus each from the spaced and massed conditions were subject to reversal. These reversed stimuli were pseudo-randomly interspersed with a non-reversed medium-reward stimulus and a non-reversed loss stimulus from each condition, yielding 8 stimuli total. In the reversal, the feedback for the first presentation of the reversed stimuli was as expected, while the remaining 9 repetitions were reversed (at a 78% probability).

We did not find any reliable effect of spaced training on reversal of reward or loss associations. Massed-trained reward-associated stimulus performance across the repetitions following the reversal (3-10) was 65.7 % [58.7 76.9]; spaced-trained performance was 60.1 % [49.1 71.1]. While performance on the spaced-trained stimuli was lower, this effect was not significant (t_(30)_ = 1.06, CI [-5.2 16.5]; p = 0.30; TOST t_(30)_ = 1.89, p = 0.034). Massed-trained loss-associated stimulus performance across the repetitions following the reversal was 36.3% [23.6 49.0]; spaced-trained performance was 38.3% [23.9 52.8] (t_(30)_ = −0.26, CI [-18.0 13.0]; p = 0.80; TOST equivalence test, t_(30)_ = −2.69, p = 0.005). As the reversal phase came after a long experiment before and during scanning, including an unrelated demanding cognitive control task, it is possible that the results were affected by general fatigue. The lack of an effect of spaced training of reversal performance indicates that alternative cognitive or short-term learning mechanisms can override well-learned reward associations.

### Three-week follow-up

We administered a follow-up test of memory for the value of conditioned stimuli approximately 3 weeks later (mean, 24.5 days; range, 20-37 days). An online questionnaire was constructed with each participant’s stimuli using Google Forms (https://docs.google.com/forms). Participants were instructed to try to remember whether a stimulus was associated with winning money or not winning money, using an adapted version of the scan from the rating phase of the in-lab experiment. Responses were recorded using a 10-point radio button scale, anchored with “0% lucky” on the left to “100% lucky” on the right. Similar to the in-lab ratings, participants were instructed to respond to the far right end of the scale if they were completely confident that a given stimulus was associated with reward and to the far left if they were completely confident that a given stimulus was associated with no reward. Thus, distance from the center origin represented confidence in their memory. Note that no choice test measures were collected at the long-term follow-up.

In-lab portions of the study were presented using Psychtoolbox 3.0 (Brainard, 1997), with the initial in-lab session conducted on 21.5” Apple iMacs. Online training was completed using expfactory.org (Sochat et al., 2016), with functions adapted from the jspsych library (de Leeuw, 2015). At the second in-lab session, before scanning, participants completed massed-stimulus training on a 15” MacBook Pro laptop. During scanning, stimuli were presented on a screen positioned above the participant’s eyes that reflected an LCD screen placed in the rear of the magnet bore. Responses during the fMRI portion were made using a 5-button cylinder button response box (Current Designs, Inc.). Participants used the top button on the side of the cylinder for “Yes” responses and the next lower button for “No” responses. We positioned the response box in the participant’s hand so that the arrangement mirrored the relative position of the up and down arrow keys on the keyboard from the training task sessions.

### Experimental design, Study 2

The procedure for Study 2 was the same as for Study 1, with the important difference that the long-term follow-up was conducted in the lab rather than online. There were two smaller differences: learning for massed stimuli was conducted in full without interruption for intermediate ratings and choices and fMRI data were not collected.

Stimuli for Study 2 were composed of the most neutrally-rated landscape stimuli from Study 1 pre-experiment ratings. Two counterbalance stimulus lists were created and assigned randomly to participants. The initial learning session for the spaced-trained stimuli and the three online training sessions were completed as described above. Following the training and testing phases, participants completed the OSPAN to collect a measure of working memory capacity. Scores were calculated as in Study 1 (mean, 49.7; range 17-83); mean performance on the arithmetic component was 93.1%.

During the two weeks between the in-lab sessions, participants completed three short “massed” online training sessions for the spaced-trained stimuli, as described above. We found that data for three sessions in one participant were missing, data for two sessions in one participant were missing, and data for one session in 5 participants was missing. Based on the information from Study 1, we can infer that some data was missing for technical reasons and some missing because of non-compliance. Among participants with at least one missing online session, performance during scanning for spaced-trained stimuli was near the group mean (84.8% vs. 86.4%). Note that the absence of spaced training in some participants would, if anything, weaken any differences between the spaced and massed condition.

### Second in-lab session

The second in-lab session was completed approximately two weeks after the first session (mean, 12.8 days; range, 10-17 days). Here, participants engaged in the initial massed learning session, which then continued through all 25 repetitions of each “massed” stimulus. Short rest breaks were included, but Study 2 omitted the intervening reward rating and choice test phases of Study 1. In the last part of the learning phase, to assess end-state performance on both spaced-trained and massed-trained stimuli, 3 repetitions of each stimulus were presented in a pseudo-random order. Rating and choice phase data were acquired after this learning block, with trial timing as described above.

After the choice phase, we administered an exploratory phase to assess potential conditioned stimulus-cued biases in new learning. This phase was conducted in a subset of 25 participants, as the task was still under development when the data from the initial 6 participants were acquired. Participants engaged in learning about new stimuli (abstract characters) in the same paradigm as described above (**Fig. 1B**) while unrelated spaced-or massed-trained landscape stimuli were presented tiled in the background during the choice period. Across all trials, we found a positive influence of background prime reward value on the rate of “Yes” responding (reward prime mean 54.4 % CI [48.2 60.4]; loss prime mean 43.2 % CI [36.6 48.0). This did not differ between the spaced and massed conditions (spaced difference, 13.0 % CI [4.4 21.6]; massed difference, 11.0 % CI [1.6 20.4]; t_(24)_ = 0.71, CI [-3.8 7.8]; p = 0.49; TOST equivalence test, p = 0.017). One limitation in this exploratory phase was that learning for the new stimuli, similar to that reported below for the regular phases, was quite rapid, likely due to the sequential ordering of the first and second presentations of a new stimulus (performance reached 77.5 % correct by the second repetition). Rapid learning about the new stimuli may have minimized the capacity to detect differences in priming due to spaced vs. massed training.

### Three-week follow-up

Approximately 3 weeks after the second in-lab session (mean, 21.1 days; range, 16-26 days), participants returned to the lab for the third and final in-lab session. Using the same testing rooms as during the previous sessions (which included the full training session on massed stimuli), participants completed another rating phase. Participants were reminded of the reward rating instructions and told to “do their best” to remember whether individual stimuli had been associated with reward or loss during training. Trial timing was as described above, and the order of stimuli was pseudo-randomized. As in Study 1, we did not collect any choice test data in the follow-up session.

### fMRI Data Acquisition

Whole-brain imaging was conducted on a GE 3T Discovery system equipped with a 32-channel head coil (Stanford Center for Cognitive and Neurobiological Imaging). Functional images were collected using a multiband (simultaneous multi-slice) acquisition sequence (TR = 680 ms, TE = 30 ms, flip angle = 53, multiband factor = 8; 2.2 mm isotropic voxel size; 64 (8 by 8) axial slices with no gap). For participant 290, TR was changed due to error, resulting in runs of 924, 874, and 720 ms TRs. Slices were tilted approximately 30° relative to the AC–PC line to improve signal-to-noise ratio in the orbitofrontal cortex (Deichmann et al., 2003). Head padding was used to minimize head motion.

During learning phase scanning, two participants was excluded for excessive head motion (5 or more >1.5 mm framewise displacement translations from TR to TR). No other participant’s motion exceeded 1.5 mm in displacement from one volume acquisition to the next. For seven other participants with 1 or more events of >0.5-mm displacement TR-to-TR, any preceding trial within 5 TRs and any current/following trial within 10 subsequent TRs of the motion event were excluded from multivariate analyses; for univariate analyses, these trials were removed from regressors of interest. For participant 310, the display screen failed in the middle of the first learning phase scanning run. This run was restarted at the point of failure and functional data were concatenated. For four participants, data from the final no-feedback fMRI block was not collected due to time constraints. Additionally, for the no-feedback block three participants were excluded for excessive head motion, leaving 26 remaining participants for the no-feedback phase analysis.

For each functional scanning run, 16 discarded volumes were collected prior to the first trial to both allow for magnetic field equilibration and to collect calibration scans for the multiband reconstruction. During the scanned learning phase, two functional runs of an average of 592 TRs (6 min and 42 s) were collected, each including 50 trials.

During the no-feedback phase, one functional runs of an average of 722 TRs (8 min and 11 s) was collected, including 160 trials. Structural images were collected either before or after the task, using a high-resolution T1-weighted magnetization prepared rapid acquisition gradient echo (MPRAGE) pulse sequence (0.9 × 0.898 × 0.898 mm voxel size).

### Behavioral analysis

Behavioral analyses were conducted in Matlab 2016a (The MathWorks, Inc., Natick, MA). Results presented below are from the following analyses: t-tests vs. chance for learning performance, within-group (paired) t-tests comparing differences in reward-and loss-associated stimuli across conditions, Pearson correlations, and Fisher z-transformations of correlation values. We additionally tested whether non-significant results were weaker than a moderate effect size using the Two One-Sided Test (TOST) procedure (Schuirmann, 1987; Lakens, 2017) and the TOSTER library in R (Lakens, 2017). We used bounds of Cohen’s *d* = 0.51 (Study 1) or *d* = 0.53 and *d* = 0.54 (Study 2), where power to detect an effect in the included group of participants is estimated to be 80%.

End-state learning accuracy in Study 1 averaged across the last 5 of 6 repetitions in the scanned intermixed learning session. End-state learning accuracy for Study 2 averaged across the last 2 of 3 repetitions in the final intermixed learning phase. For the purpose of correlations with working memory, initial learning repetitions 2-10 were averaged (as repetition 1 cannot reflect learning). In Study 1, the post-learning ratings were taken from the ratings collected before the scan (after 25 repetitions across all massed-and spaced-trained stimuli). In Study 2, the post-learning ratings were collected after all learning repetitions were completed.

For the analysis of maintenance of learned values in Study 2, we computed a percentage maintenance measure. This was calculated by dividing the long-term reward rating difference (reward-minus loss-associated mean stimulus ratings) by the post-learning rating difference. The same analysis but with a range restricted to a minimum of 0 (eliminating any reversals in ratings) and a maximum of 100% yielded similar results but with lower variance and correspondingly higher t-value.

### fMRI Data Analysis

Data from all participants were preprocessed several times to fine tune the parameters. After each iteration the decision to modify the preprocessing was purely based on the visual evaluation of the preprocessed data and not based on results of model fitting. Results included in this manuscript come from application of a standard preprocessing pipeline using FMRIPREP version 1.0.0-rc2 (http://fmriprep.readthedocs.io), which is based on Nipype (Gorgolewski et al., 2011). Slice timing correction was disabled due to short TR of the input data. Each T1 weighted volume was corrected for bias field using N4BiasFieldCorrection v2.1.0 (Tustison et al., 2010), skullstripped using antsBrainExtraction.sh v2.1.0 (using the OASIS template), and coregistered to skullstripped ICBM 152 Nonlinear Asymmetrical template version 2009c (Fonov et al., 2009) using nonlinear transformation implemented in ANTs v2.1.0 (Avants et al., 2008). Cortical surface was estimated using FreeSurfer v6.0.0 (Dale et al., 1999).

Functional data for each run was motion corrected using MCFLIRT v5.0.9 (Jenkinson et al., 2002). Distortion correction for most participants was performed using an implementation of the TOPUP technique (Andersson et al., 2003) using 3dQwarp v16.2.07 distributed as part of AFNI (Cox, 1996). In case of data from participants 8, 12, 14, 27, and 36 spiral fieldmaps were used to correct for distortions due to artifacts induced by the TOPUP approach in those participants. This decision was made based on visual inspection of the preprocessed data prior to fitting any models. The spiral fieldmaps were processed using FUGUE v5.0.9 (Jenkinson, 2003). Functional data was coregistered to the corresponding T1 weighted volume using boundary based registration 9 degrees of freedom - implemented in FreeSurfer v6.0.0 (Greve and Fischl, 2009). Motion correcting transformations, field distortion correcting warp, T1 weighted transformation and MNI template warp were applied in a single step using antsApplyTransformations v2.1.0 with Lanczos interpolation. Framewise displacement (Power et al., 2014) was calculated for each functional run using Nipype implementation. For more details of the pipeline see http://fmriprep.readthedocs.io/en/1.0.0-rc2/workflows.html.

General linear model analyses were conducted using SPM (SPM12; Wellcome Trust Centre for Neuroimaging). MRI model regressors were convolved with the canonical hemodynamic response function and entered into a general linear model (GLM) of each participant’s fMRI data. Six scan-to-scan motion parameters (x, y, z dimensions as well as roll, pitch, and yaw) produced during realignment were included as additional regressors in the GLM to account for residual effects of participant movement.

We first conducted univariate analyses to identify main effects of value and reward in the learning phase, as well as effects of presentation without feedback in the final phase. The learning phase GLM included regressors for the stimulus onset (2 s duration) and feedback onset (2 s duration). The stimulus onset regressor was accompanied by a modulatory regressor for reward value (reward vs. loss), separately for spaced-and massed-trained stimuli. The feedback regressor was accompanied by four modulatory regressors for reward value (reward vs. loss) and spacing (spaced-vs. massed-trained). The median performance in the scanner was 97.5%, and because learning was effectively no longer occurring during the scanning phase, we did not use a reinforcement learning model to create regressors.

The no-feedback phase GLM included regressors for the stimulus onset (1.5 s duration) and query onset (3.0 s duration). In the no-feedback phase, we conducted an exploratory cross-stimulus repetition-suppression analyses (XSS; Klein-Flugge et al., 2013). Here, non-perceptual features associated with a stimulus are predicted to activate the same neural population representing the feature. This feature coding is then predicted to lead to a suppressed response in subsequent activations, for example, when a different stimulus sharing that feature is presented immediately after the first stimulus (Barron et al., 2016). In the XSS model, we contrasted sequential presentations of stimuli that shared value association (reward and loss) and spacing (spaced vs. massed), yielding four regressors. For example, if two different reward-associated and spaced-trained stimuli followed in successive trials, the first trial would receive a 1 value and the second trial would receive a −1. These regressors were entered into contrasts to yield reward vs. non XSS for spaced-trained stimuli and reward vs. non XSS for massed-trained stimuli.

For multivariate classification analyses, we estimated a mass-univariate GLM where each trial was modeled with a single regressor, giving 100 regressors for the learning phase. The learning phase regressor duration modeled the 2 seconds long initial stimulus presentation period. Models included the 6 motion regressors and block regressors as effects of no interest. Multivariate analyses were conducting using The Decoding Toolbox (Hebart et al., 2014). Classification utilized a L2-norm learning support vector machine (LIBSVM; Chang and Lin, 2011) with a fixed cost of *c* = 1. The classifier was trained on the full learning phase data, with the two scanning blocks subdivided into four runs (balancing the number of events within and across runs). We conducted four classification analyses: overall reward-vs. loss-associated stimulus classification, spaced-vs. massed-trained stimulus classification, and reward-vs. loss-associated stimulus classification separately for spaced-and massed-trained stimuli. For the final two analyses, the results were compared to test differences in value classification performance for spaced vs. massed stimuli. Leave-one-run-out cross-validation was used, with results reported in terms of percent correct classification. Statistical comparisons were made using t-tests vs. chance (50%); for the comparison of two classifier results, paired t-tests were used.

In addition to the two ROI analyses, we conducted a searchlight analysis using The Decoding Toolbox (Hebart et al., 2014). We used a 4-voxel radius spherical searchlight. Training of the classifier and testing were conducted as described above for the region of interest MVPA. Individual subject classification accuracy maps were smoothed with a 4mm FWHM kernel prior to group-level analysis. A comparison between value classification between spaced-and massed-trained stimuli was conducted using a t-test on the difference between participant’s spaced-and massed-trained classification SPMs (equivalent to a paired t-test).

For both univariate and searchlight results, linear contrasts of univariate SPMs were taken to a group-level (random-effects) analysis. We report results corrected for family-wise error (FWE) due to multiple comparisons (Friston et al., 1993). We conduct this correction at the peak level within small volume ROIs for which we had an a priori hypothesis or at the whole-brain cluster level (in each case using a cluster-forming threshold of p < 0.005 uncorrected). The striatum and MTL (including hippocampus and parahippocampal cortex) ROIs were adapted from the AAL atlas (Tzourio-Mazoyer et al., 2002). The striatal mask included the caudate and putamen, as well as the addition of a hand-drawn nucleus accumbens mask (Wimmer et al., 2012). All voxel locations are reported in MNI coordinates, and results are displayed overlaid on the average of all participants’ normalized high-resolution structural images using xjview and AFNI (Cox, 1996).

## Data availability

Complete behavioral data are publicly available on the Open Science Framework (www.osf.io/z2gwf/). Unthresholded whole-brain fMRI results are available on NeuroVault (https://neurovault.org/collections/3340/) and the full fMRI dataset is publicly available on OpenNeuro (https://openneuro.org/datasets/ds001393/versions/00001).

## Results

Across two studies, we measured learning and maintenance of conditioned stimulus-value associations over time. In the first in-lab session, participants learned stimulus-value associations for a set of “spaced-trained” stimuli (the spaced initial learning session). Over the course of the next two weeks, participants engaged in three short “massed” training sessions online (the spaced online training sessions). Participants then returned to complete a second in-lab session, where they learned stimulus-value associations for a new set of “massed-trained” stimuli (the massed initial learning session and continued training). All learning for the massed-trained stimuli occurred consecutively in the same session. By the end of training on the massed-trained stimuli, experience was equated between the spaced-and massed-trained stimuli. While the timing of trials was equivalent across the spaced-trained and massed-trained stimuli, the critical difference was that multiple days were inserted in-between the short training sessions for spaced-trained stimuli. 3-weeks after the second in-lab session, participants completed a long-term follow-up reward rating measure.

### Study 1

#### Learning of value associations

Participants rapidly acquired the best “Yes” or “No” response for the reward-or loss-associated stimuli during the initial spaced (lab session 1) and massed (lab session 2) learning sessions. By the second repetition of each stimulus, accuracy quickly increased to 89.1 % (95% Confidence Interval (CI) [87.4 95.2]) for spaced-trained stimuli and 91.3 % (CI [84.7 93.5]) for massed-trained stimuli (p-values < 0.001). Participants exhibited a noted bias (77.7 %) toward “Yes” responses for the first trial of a given stimulus when no previous information could be used to guide their response. By the end of the initial learning sessions (repetition 10), performance increased to 83.3 % (CI [76.8 89.8]) for the spaced-trained stimuli and 93.6 % (CI [90.4 96.7]) for the massed-trained stimuli (**Fig. 2A**). Performance was higher by the end of the initial learning session for the massed-trained stimuli (t_(32)_ = 3.13, CI [3.6 17.0]; p = 0.0037). Note that the only difference between the spaced and massed learning sessions is that there is greater task exposure at the time of the massed learning session; both sessions have the same within-session trial timing and spacing. After the completion of the online learning sessions for spaced-trained stimuli and further in-lab learning for massed-trained stimuli, as expected, we found that participants showed no significant difference in performance across conditions (repetitions 27-31; spaced-trained, 92.1 % CI [88.3 96.0]; massed-trained, 94.7 % CI [91.9 97.6]; t_(32)_ = 1.59, CI [-1.0 5.9]; p = 0.123; **Fig. 2A**). However, this effect was not statistically equivalent to a null effect, as indicated by an equivalence test using the TOST procedure (Lakens, 2017): the effect was not significantly within the bounds of a medium effect of interest (Cohen’s d = ± 0.51, providing 80% power with 33 participants; t_(32)_ = 1.34, p = 0.094), and thus we cannot reject the presence of a medium-size effect.

**Figure 2.**
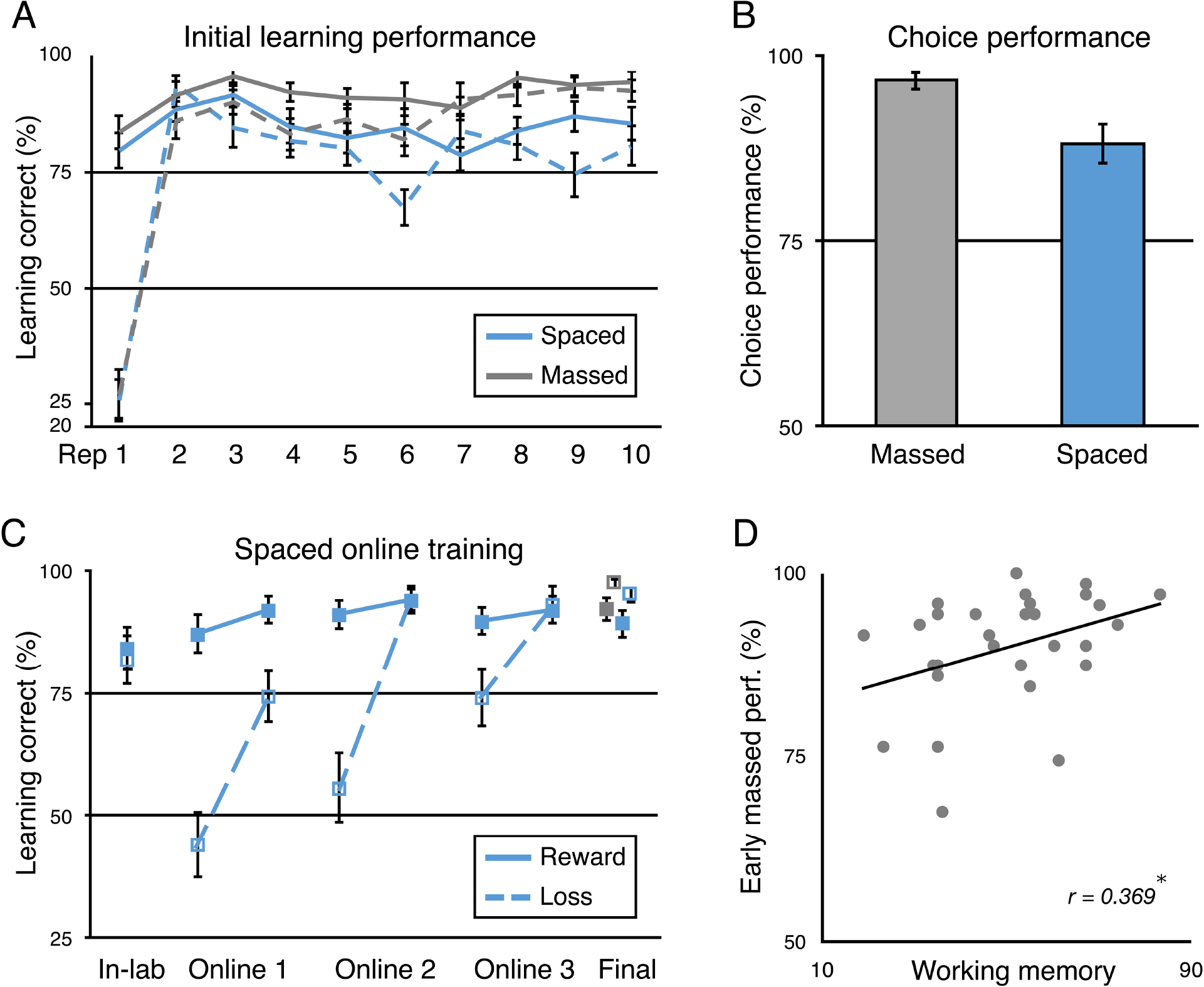
Study 1 learning results. ***A***, Performance in the initial learning sessions for the spaced-and massed-trained stimuli across the first ten repetitions of each stimulus. (Massed stimuli in grey; spaced in blue; reward-associated stimuli in solid lines; loss-associated stimuli in dotted lines.) ***B***, Incentivized two-alternative forced-choice performance between reward-and loss-associated stimuli following the initial spaced and massed learning sessions. ***C***, Spaced performance across online learning sessions and terminal performance for spaced and massed stimuli. Performance is depicted for the last in-lab repetition and the first and last (fifth) repetition of each stimulus per online session, followed by the average of the final 27-31 repetitions in the second in-lab session including fMRI. ***D***, Positive correlation between early massed-trained stimulus learning phase performance and working memory capacity (O-SPAN). (* p < 0.05). Rep. = repetition. Error bars represent one standard error of the mean (s.e.m.).

Performance in the initial session illustrated that participants learned the reward value of the stimuli during learning. First, participants showed higher learning accuracy for high reward (mean $0.45 feedback) vs. medium reward (mean $0.25) spaced-trained stimuli in the second half of learning (high reward, 90.0 % CI [83.2 96.8]; medium reward 78.0 % CI [70.7 85.4]; t_(32)_ = 2.98, CI [3.8 20.1]; p = 0.0054). After extensive training in the task, however, we did not observe a similar effect for initial learning of the massed-trained stimuli (high reward, 91.9 % CI [85.8 97.9]; medium reward, 93.2 % CI [88.5 98.0]; t_(32)_ = −0.38, CI [-8.6 5.9]; p > 0.70; TOST = t_(32)_ = 2.54, p = 0.008). Second, after the initial learning phase participants completed an incentivized two-alternative forced choice test phase. Here, no trial-by-trial feedback was given, but additional rewards were paid based on performance. Participants exhibited a strong preference for the reward-vs. loss-associated stimuli in choices between both spaced-and massed-trained stimuli (spaced accuracy, 96.5 % CI [82.6 93.6]; massed accuracy, 88.1 % CI [94.4 98.8]; p-values < 0.00001; **Fig. 2B**).

After the first in-lab session, participants continued learning about the set of spaced-trained stimuli across three short “massed” online sessions. We found that across the 3 online sessions, mean performance increased for loss-associated stimuli (one-way ANOVA; F_(2,72)_ = 9.26, p = 0.003; **Fig. 2B**) but not for reward-associated stimuli (F_(2,72)_ = 0.53, p = 0.59). This increase in performance for loss-associated stimuli was accompanied by a significant decrease in performance between sessions (mean change from end of session to beginning of next session: t_(24)_ = 4.71, CI [14.5 37.1]; p < 0.001) but not for reward-associated stimuli (t_(24)_ = 0.38, CI [-3.4 5.0]; p = 0.704). Performance at the beginning of the online sessions may have been influenced by a response bias toward “Yes”, as also shown in first responses to stimuli in initial learning (**Fig. 2A**). Forgetting that leads to a bias under uncertainty would decrease memory performance for loss-associated stimuli. However, a bias would mask any forgetting for reward-associated stimuli, as it would lead to higher performance. Thus, we cannot rule out the forgetting of reward-associated stimuli in the current design.

During the second in-lab (fMRI) session, learning performance was above 90% for both conditions, but massed-trained stimuli showed higher performance than spaced-trained stimuli (spaced choice performance, scan repetitions 2 to 6, 92.1 % CI [88.3 96.0]; massed, 94.7 % CI [91.9 97.5]; p = 0.01; CI [3.0 20.6]; t(32) = 2.74; **Fig. 2C**).

After sufficient general experience in the task, we expected to find a positive relationship between learning performance for new stimuli and working memory. We thus estimated the correlation between learning during the initial acquisition of massed-trained stimulus-value associations during the second in-lab session with the operations span measure of working memory. We found that learning performance on the massed-trained stimuli positively related to working memory capacity (r = 0.369, p = 0.049; **Fig. 2C**). Initial performance for spaced-trained stimuli did not correlate with working memory (r = −0.097, p = 0.617; TOST equivalence test providing 80% power in range r ± 0.34, p = 0.080, and thus we cannot reject the presence of a medium-size effect). The correlation between working memory and massed performance was significantly greater than the correlation with spaced performance (z = 2.16, p = 0.031). In contrast to the predicted effect for massed performance in the second session, we did not predict a relationship between first session spaced condition performance and working memory. While working memory clearly contributed to spaced learning performance, given the rapid shift in responding to loss-associated stimuli after the first trial (**Fig. 2A**), absent a prolonged practice session, working memory is also likely to be utilized to maintain task instructions (Cole et al., 2013). Initial task performance is also likely to be affected by numerous other noise-introducing factors such as the acquisition of general task rules (“task set”) and adaptation to the testing environment. However, the lack of a correlation with working memory in the spaced may also indicate that the working memory correlations in general are weak and hard to detect if present, even with more than 30 participants. Future studies are needed to further investigate the effects of working memory on initial learning and task acquisition. When interpreting these working memory correlations with respect to previous studies on the contribution of working memory to feedback-based learning (Collins and Frank, 2012), it is important to note that the 8 stimuli in the spaced and massed condition were introduced in two sequential sets of 4 stimuli. Thus, participants would only need to maintain 4 instead of 8 stimulus-reward or stimulus-response in short-term memory, well within the range reported in previous studies.

#### Long-term maintenance

Next, we turned to the critical question of whether spaced training over weeks led to differences in long-term memory for conditioned reward associations. Baseline reward ratings were collected before the fMRI scanning in the second in-lab session. Higher ratings indicate strong confidence in a reward association while lower ratings indicate higher confidence in a neutral/loss association; ratings more toward the middle of the scale indicated less confidence (**Fig. 1C**). After training but before fMRI scanning, when experience was matched across the spaced and massed conditions, we found that ratings across condition clearly discriminated between reward-and loss-associated stimuli (spaced rating difference, 47.5 % CI [39.2 55.7]; massed rating difference, 62.5 % CI [56.5 68.4]; p-values < 0.00001; condition difference, t_(32)_ = 2.73, CI [3.0 20.6]; p = 0.01; **Fig. 3A**, left).

**Figure 3.**
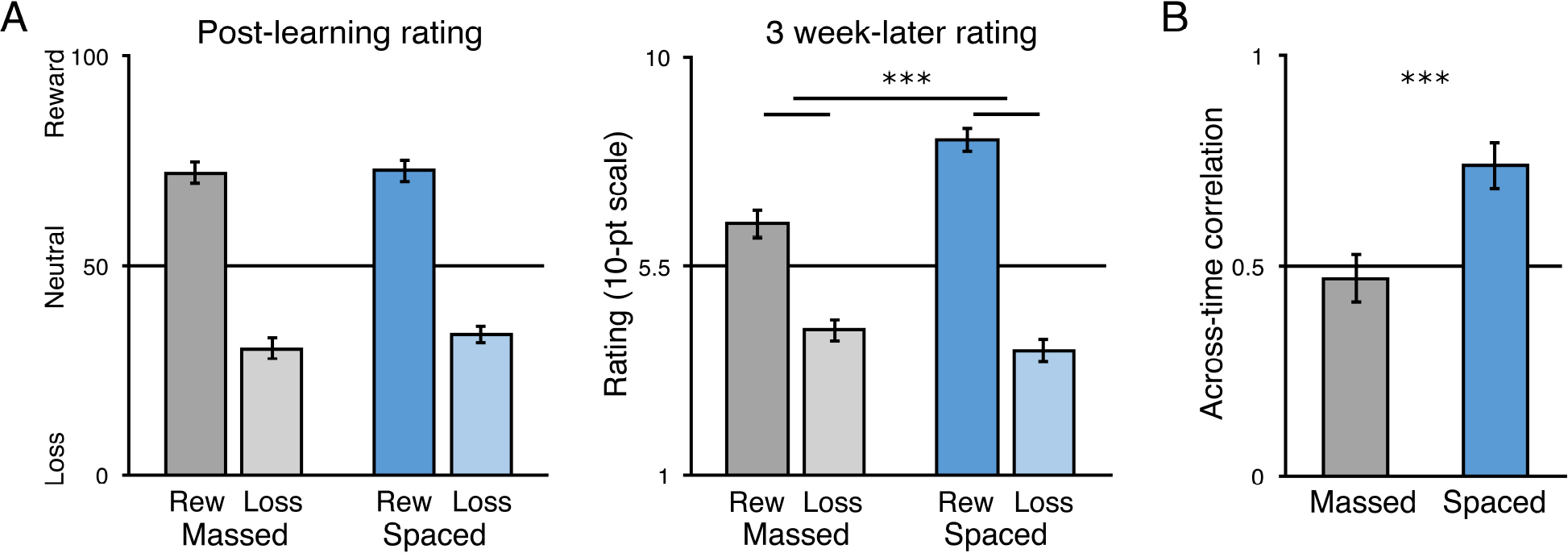
Study 1 post-learning value association strength and long-term maintenance of value associations. ***A***, Post-learning reward association ratings for the massed-and spaced-trained stimuli (left); 3-week-later reward association ratings (right). Reward-associated stimuli in darker colors; loss-associated stimuli in lighter colors. ***B***, Average of the correlation (r) within-participant of massed-trained stimulus reward ratings and spaced-trained stimulus reward-ratings (statistics were computed on z-transformed ratings). (*** p < 0.001). Error bars, s.e.m.

At the long-term follow-up, only rating data were collected. Importantly, to validate the use of the reward rating scale in the follow-up measures, we tested how strongly these measures were related. We found that within-participants, massed-trained ratings were strongly correlated with preferences for stimuli in the separate choice test phase (mean r = 0.92, CI [0.88 0.95]; range 0.68-1.00; t-test on z-transformed r-values, t_(32)_ = 11.10 CI [1.79 2.59]; p < 0.0001). This strong correlation indicates that the reward ratings capture the same underlying values learned via feedback learning as the forced-choice test measure commonly used as an assessment of learning.

To measure long-term maintenance of conditioning, after approximately 3 weeks, participants completed an online questionnaire on reward association strength using a 10-point scale. The instructions for ratings were the same as the in-lab ratings phase. Critically, we found that while the reward value discrimination was significant in both conditions (spaced difference, 4.55 CI [3.75 5.34]; t_(32)_ = 11.61, p < 0.001; massed difference, 2.24 CI [1.59 3.01]; t_(32)_ = 6.60, p < 0.001), reward value discrimination was significantly stronger in the spaced than in the massed condition (t_(32)_ = 4.55, CI [1.23 3.25]; p < 0.001; **Fig. 3B**). This effect was driven by greater maintenance of the values of reward-associated stimuli (spaced vs. massed, t_(32)_ = 4.73, CI [1.04 2.58]; p < 0.001; loss spaced vs. massed, t_(32)_ = −1.37, CI [-1.08 0.21]; p = 0.18; TOST equivalence test, t_(32)_ = 1.56, p = 0.064, n.s.). Note that the benefit of spacing at the long-term follow-up also differs from the baseline at the end of learning, where performance was marginally higher for massed-trained stimuli.

Next, we analyzed the consistency of ratings from the end of learning to the long-term follow-up. The post-learning ratings were collected on a graded scale and the 3-week follow-up rating was collected on a 10-point scale; this prevents a direct numeric comparison but allows for a correlation analysis. Such an analysis can test whether ratings in the massed case were simply scaled down (preserving ordering) or if actual forgetting introduced noise (disrupting an across-time correlation). We predicted that the value association memory for massed-trained stimuli actually decayed, leading to a higher correlation across time for spaced-trained stimuli. We indeed found that ratings were significantly more correlated across time in the spaced-trained condition (spaced r = 0.74, CI [0.63 0.85]; massed r= 0.47, CI [0.35 0.59]; t-test on z-transformed values, t_(32)_ = 4.13, CI [1.28 0.44]; p < 0.001). While the correlation for the spaced-trained stimuli was high (median r = 0.85), there was still variability in group, with individual participant r-values ranging from −0.36 to 1.0. Overall, these results indicate that spaced-trained stimuli exhibited significantly stronger long-term memory for conditioned associations and more stable memory than massed-trained stimuli.

One limitation to these results is that in the current design, cues in the learning environment may bias performance in favor of the spaced-trained stimuli: online training for spaced stimuli was conducted outside the lab, likely on the participant’s own computer, which was likely the same environment for the 3-week follow-up measure. While it seems unlikely that a testing environment effect would fully account for the large difference in long-term maintenance that we observed, we conducted a second study to replicate these results in a design where the testing conditions would if anything bias performance in favor of the massed-trained stimuli.

### Study 2

#### Learning of value associations

In Study 2, our aim was to replicate the findings of Study 1 and to extend them by conducting the 3-week follow-up session in the lab, allowing for a direct comparison with post-learning performance. Learning sessions for spaced-and massed-trained stimuli were the same as in Study 1, with the exception that massed learning in Study 2 omitted the mid-learning assessment with ratings and choices. During the initial spaced and massed learning sessions, by the second trial, accuracy had increased to 89.9 % (CI [85.3 94.6]) for spaced-trained stimuli and to 87.2 % (CI [84.1 93.3]) for massed-trained stimuli (p-values < 0.001). As before, participants exhibited a noted bias (67.2 %) toward “Yes” responses for the first trial of a given stimulus when no previous information could be used to guide their response. By the end of the initial learning sessions, performance was at a level of 84.3 % (CI [79.3 89.3]) for the spaced-trained stimuli and 86.2 % (CI [81.2 91.1]) for the massed-trained stimuli (**Fig. 4A**), which was matched across conditions (10^th^ repetition; t_(30)_ = 0.59, CI [-4.72 8.52]; p = 0.56; TOST equivalence test within a range of Cohen’s d = ± 0.53, providing 80% power with 31 participants; t_(30)_ = 2.37, p = 0.012). By the end of training, after the online sessions for spaced-trained stimuli and the completion of the in-lab learning for massed-trained stimuli, we found that performance was equivalent across conditions (spaced-trained, 86.4 % CI [82.2 90.6]; massed-trained, 87.1 % CI [81.4 92.8]; t_(30)_ = 0.248, CI [-4.76 6.08]; p = 0.806; TOST equivalence test, t_(30)_ = 2.70, p = 0.006; **Fig. 4C**).

**Figure 4.**
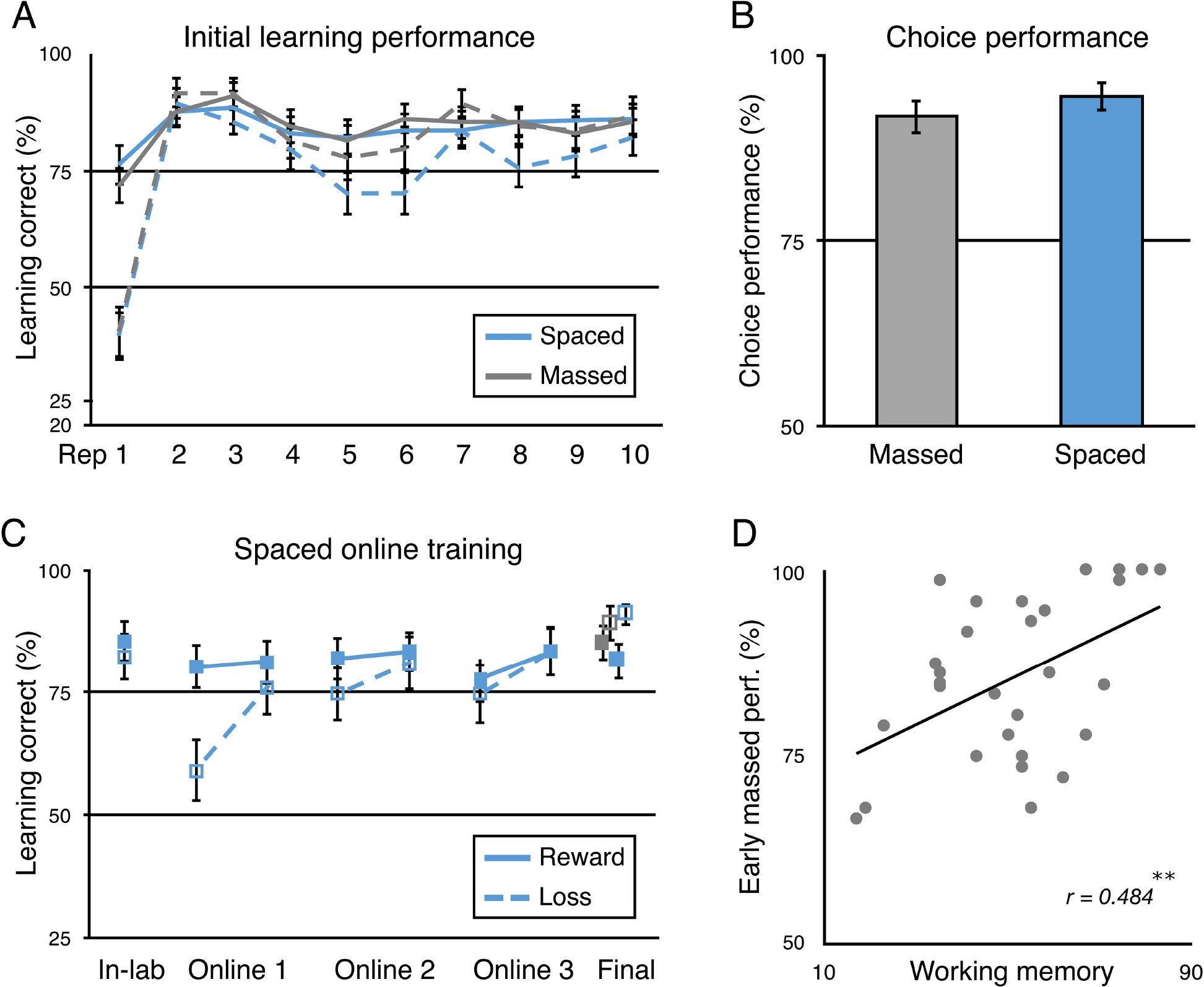
Study 2 learning results. ***A***, Performance in the initial spaced and massed learning sessions across the first ten repetitions of each stimulus. (Massed stimuli in grey; spaced in blue; reward-associated stimuli in solid lines; loss-associated stimuli in dotted lines.) ***B***, Incentivized two-alternative forced-choice performance between reward-and loss-associated stimuli following the completion of all learning repetitions. ***C***, Spaced performance across training and terminal performance for spaced and massed stimuli. Performance is shown for the last in-lab repetition and the first and last (fifth) repetition of each stimulus per online session. Terminal performance is represented as the average of the final 2 repetitions of each stimulus in the last learning session. ***D***, Positive correlation between early massed-trained stimulus learning performance and working memory capacity (OSPAN). (** p < 0.01). Error bars, s.e.m.

As in Study 1, performance in the initial learning sessions illustrates that participants learned the reward value of the stimuli during learning. First, participants tended to prefer the high reward vs. medium reward spaced-trained stimuli during the second half of learning (high reward, 90.3 % CI [84.7 96.0]; medium reward, 79.9 % CI [71.0 88.7]; t_(30)_ = 2.03, CI [0.0 20.9]; p = 0.051). Later, however, after extensive training in the task, we did not observe a similar effect for initial learning of the massed-trained stimuli (high reward, 84.8 % CI [75.5 94.2]; medium reward, 85.5 % CI [77.7 93.3]; t_(30)_ = −0.11, CI [-12.5 11.2]; p > 0.91; TOST = t_(30)_ = 2.84, p = 0.004). Second, performance in the incentivized forced-choice test phase after the completion of learning showed strong preference for the reward-vs. loss-associated stimuli in choices between both spaced-and massed-trained stimuli (spaced-trained, 94.6 % CI [87.2 96.3]; massed-trained, 91.7 % CI [90.8 98.3]; difference between conditions, p > 0.26; TOST equivalence test, t_(30)_ = 1.83, p = 0.04; **Fig. 4B**). Equivalent choice performance after learning for spaced-and massed-trained stimuli is important for the long-term follow-up measure.

After the initial spaced learning session in the first in-lab visit, participants continued learning about the set of spaced-trained stimuli across three short “massed” online sessions. As in Study 1, we found that across the 3 online sessions, mean performance did not change for reward-associated stimuli (one-way ANOVA; F_(2,69)_ = 0.06, p = 0.94; **Fig. 4B**). In contrast to the previous study, we did not find an increase in performance across sessions for loss-associated stimuli (F_(2,69)_ = 1.09, p = 0.34), although a post-hoc comparison of the first to the third session showed an increase (t_(23)_ = 2.43, CI [1.1 3.4]; p = 0.024). However, we did replicate the finding that loss-associated stimuli showed a significant decrease in performance between sessions (mean change from end of session to beginning of next session: t_(23)_ = 2.69, CI [2.4 18.2]; p = 0.013; reward-associated stimuli (t_(23)_ = 1.40, CI [-1.6 8.3]; p = 0.18). As discussed above, this decrease in performance evident for loss-associated stimuli could indicate forgetting of values and a return toward a default “Yes” response bias (as seen in first exposure responses; **Fig. 2A**). Such a bias would make it difficult in the current design to determine whether memories for the value of reward-associated stimuli also decayed.

As in Study 1, after sufficient general experience in the reward association learning task, we expected to find a positive relationship between performance on the reward association learning task and working memory. Indeed, we found a significant correlation between massed-stimulus performance and working memory capacity (r = 0.484, p = 0.0058; **Fig. 4C**). Initial learning performance was relatively lower in Study 2 than in Study 1, which may have helped reveal a numerically stronger correlation between massed-trained stimulus performance and working memory. Meanwhile, the relationship between working memory and initial performance for spaced-trained stimuli was weak (r = 0.040, p = 0.83; TOST equivalence test, p = 0.043, providing 80% power in range r ± 0.35; difference between massed and spaced correlation, z = 1.40 p = 0.16), as expected, given the other noise-introducing factors in initial learning performance discussed above. As in Study 1, however, working memory clearly also contributed to spaced learning performance, as demonstrated by the immediate shift in mean response to loss-associated stimuli from “Yes” to “No” after initial negative feedback (**Fig. 2A**).

#### Long-term maintenance

Next, we turned to the critical question of whether spaced training over weeks led to differences in long-term memory for conditioned reward associations. For the baseline post-learning measurement for spaced-and massed-trained stimuli, ratings were collected at the end of the complete massed-stimulus training session (**Fig. 5A**, left). Reward ratings showed strong discrimination of value (spaced-trained reward minus loss rating difference, 47.1 % CI [40.9 53.2]; massed-trained rating difference, 52.5 % CI [46.7 58.3]; condition difference, t_(30)_ = −1.86, CI [-0.5 11.4]; p = 0.073; **Fig. 5A**, left). Note that as in the previous study, we collected ratings data but no choice test data in the long-term follow-up. To again validate the use of the reward rating scale in the follow-up measures, we tested whether within-participant reward ratings were related to choice test preferences. Again, we found that ratings positively correlated with choice preference across all stimuli (mean r = 0.87, CI [0.82 0.91]; range 0.56-1.00; t-test on z-transformed r-values, t_(30)_ = 14.08, CI [1.33 1.78]; p < 0.0001). By replicating the strong correlation found in Study 1, these results indicate that reward ratings capture the essential underlying values revealed through forced-choice preferences.

**Figure 5.**
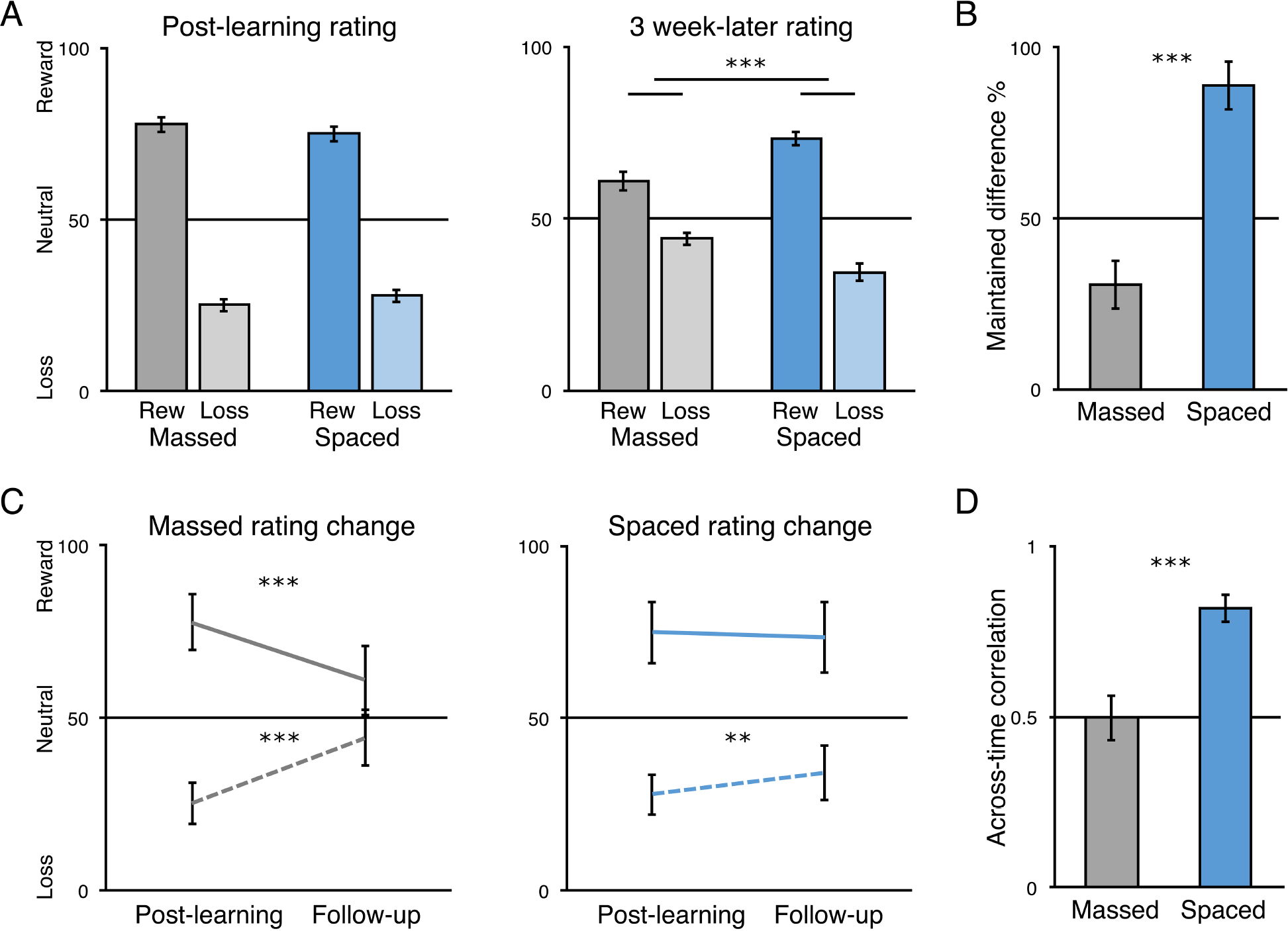
Study 2 post-learning reward association strength and maintenance of value associations. ***A***, Reward association ratings for the massed-and spaced-trained stimuli after the second in-lab session (left), and after the 3-week-later in-lab final reward association rating session (right). Reward-associated stimuli in darker colors; loss-associated stimuli in lighter colors. ***B***, Percent of initial reward association difference (reward minus loss associated rating) after the second in-lab session maintained across the 3-week delay to the third in-lab session, separately for massed-and spaced-trained stimuli. ***C***, Post-learning and 3-week follow-up ratings re-plotted within condition for reward-associated (solid line) and loss-associated stimuli (dotted line). ***D***, Average of the correlation (r) within-participant of massed-trained stimulus reward ratings and spaced-trained stimulus reward-ratings (statistics were computed on z-transformed ratings). (** p < 0.01, *** p < 0.001) Error bars, s.e.m. (A, B, D), and within-participants s.e.m (C).

To measure long-term maintenance of conditioning, after approximately 3 weeks, participants returned for a third in-lab session for a brief session where they gave reward ratings for all stimuli. Rating discrimination between reward-and loss-associated stimuli was significant in both conditions (spaced difference, 39.1 % CI [32.4 45.8]; t_(29)_ = 11.96, p < 0.001; massed difference, 16.7 % CI [9.4 24.1]; t_(29)_ = 4.65, p < 0.001). Importantly, reward value discrimination was significantly stronger in the spaced than in the massed condition (t_(29)_ = 4.98, CI [13.2 31.5]; p < 0.001; **Fig. 5A, right**). At follow-up, this stronger maintenance of learned value associations in the spaced condition was significant for both reward and loss stimuli (reward, t_(29)_ = 3.43, CI [5.0 20.0]; p = 0.0018; loss, t_(29)_ = −4.11, CI [-14.7 −5.0]; p < 0.001). The design of Study 2 allowed us to directly compare post-learning ratings and 3-week later ratings to calculate the degree of maintenance of conditioning. As expected, the difference in maintenance for reward associations was significantly greater for spaced-than massed-trained stimuli (spaced, 87.3 % CI [73.2 101.5]; massed, 30.0 % CI [16.2 43.9]; t_(29)_ = 5.49, CI [36.0 78.6]; **Fig. 5B**). Moreover, we found that ratings significantly decayed toward neutral for both reward-and loss-associated massed-trained stimuli (massed reward, t_(29)_ = −6.09, CI [-21.7 −10.8]; p < 0.001; loss, t_(29)_ = 9.95, CI [15.3 23.3]; p < 0.001). For spaced-trained stimuli, we found no decay for reward-associated stimuli but some decay for loss-associated stimuli (spaced reward, t_(29)_ = −1.21, CI [-4.0 1.0]; p = 0.23; TOST equivalence test, t_(29)_ = 1.74, p = 0.045; loss, t_(29)_ = 3.00, CI [2.1 11.4]; p = 0.0055). Interestingly, we found that the ratings for loss-associated stimuli decayed significantly more than those for reward-associated stimuli (t_(29)_ = −2.18, CI [-10.20 −0.33]; p = 0.037), an effect in line with the between-sessions drop in performance for loss-associated stimuli. We did not find a difference in ratings decay for the massed-trained stimuli (t_(29)_ = −1.05, CI [-9.01 2.89]; p = 0.302); however, this null finding could be due to floor effects, as ratings are near 50%.

Finally, as in Study 1, we predicted that the value association memory for massed-trained stimuli was not decreased by scaling but actually decayed, which would lead to a lower across-time correlation in ratings. To test this, we correlated ratings in the second in-lab session with ratings in the third in-lab session separately for massed-and spaced-trained stimuli. We replicated the finding that ratings were significantly more correlated across time in the spaced-trained condition (spaced r = 0.82, CI [0.74 0.90]; massed r = 0.50, CI [0.37 0.63]; t-test on z-transformed values, t_(29)_ = 5.22, CI [0.45 1.03]; p < 0.001).

By collecting the long-term follow-up ratings in the same lab environment as the massed training sessions, our design would, if anything, be biased to find stronger maintenance for massed-trained stimuli because the training and testing environments overlap. However, we found similar differences in long-term conditioning across Study 1 and Study 2, suggesting that testing environment was not a significant factor in our measure of conditioning maintenance. While it will be important in the future to also replicate these results in a choice situation such as a stable bandit task, the replication and extension of the findings of Study 1 provide strong evidence that spaced training leads to more robust maintenance of conditioned value associations at a delay, while performance in short-term learning is partly explained by working memory.

## fMRI Results

In Study 1, after the completion of matched training for the massed-trained associations in the second in-lab session, we collected fMRI data during an additional learning phase, where massed-and spaced-trained stimuli were intermixed. As noted above, during fMRI scanning, we found overall performance above 90 %, but a slight benefit for massed-trained stimuli (**Fig. 2C**).

Initial univariate analyses we did not reveal any value or reward-related differences in striatal or MTL responses due to spaced training (see **Table 1-1**). At stimulus onset, across conditions a contrast of reward vs. loss-associated stimuli revealed activation in the bilateral occipital cortex and right somatomotor cortex (whole-brain FWE-corrected p < 0.05; **Table 1-1**; unthresholded map available at https://neurovault.org/images/63125/), with no differences due to spaced-vs. massed-trained stimuli. At feedback, we found expected effects of reward (hit) vs. non-reward (miss) feedback for reward-associated stimuli in the ventral striatum (x, y, z: −10, 9, −8; z = 4.48, p = 0.019 whole-brain FWE-corrected) and VMPFC (−15, 51, −1; z = 4.93, p < 0.001 FWE; see **Table 1-1** and https://neurovault.org/images/59042/). Across conditions, loss (miss) vs. neutral (hit) feedback activated the bilateral anterior insula and anterior cingulate (**Table 1-1**; https://neurovault.org/images/63127/). However, we found that miss versus hit feedback elicited greater responses in the anterior insula and anterior cingulate cortex for loss-associated stimuli than for reward-associated stimuli (**Table 1-1**; **Fig. 6-1**; https://neurovault.org/images/63126/). Loss feedback led to greater activity for massed-vs. spaced-trained stimuli in the bilateral DLPFC, parietal cortex, and ventral occipital cortex (**Table 1-1**). A second model contrasting spaced-vs. massed-trained stimuli across value revealed no significant differences in subcortical regions of interest or in the whole brain. In a subsequent no-feedback scanning block, we examined the effect of cross-stimulus repetition-suppression (XSS) for reward-vs. loss-associated stimuli. We found no differences due to condition, but several clusters that showed overall repetition-enhancement by value, including the right dorsolateral PFC and anterior insula (**Table 1-1**). While our univariate results exhibited no clear differences based on spacing condition, they do align well with previous results on reward-based learning in human fMRI studies (Bartra et al., 2013).

**Table 1-1.**
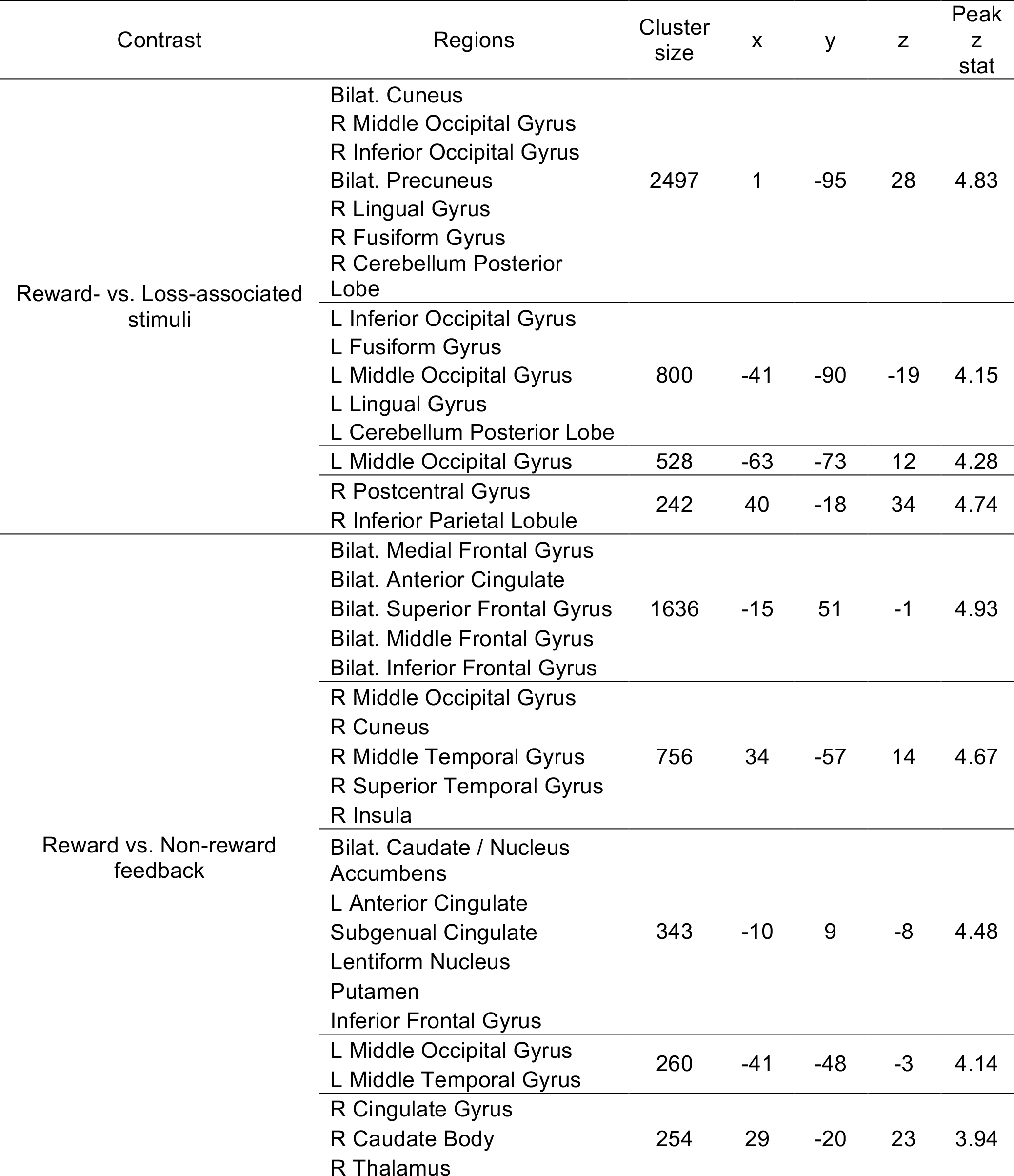
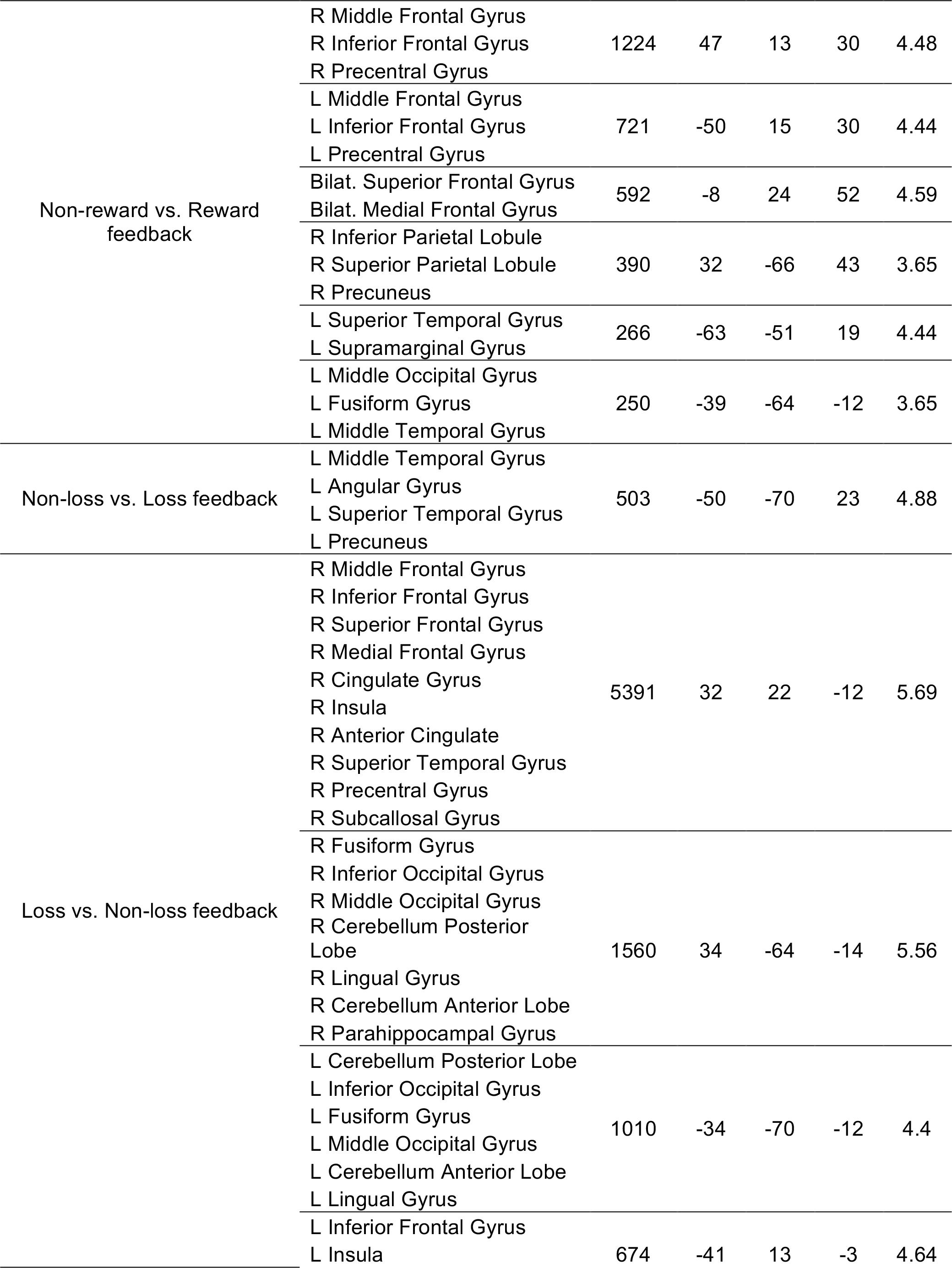
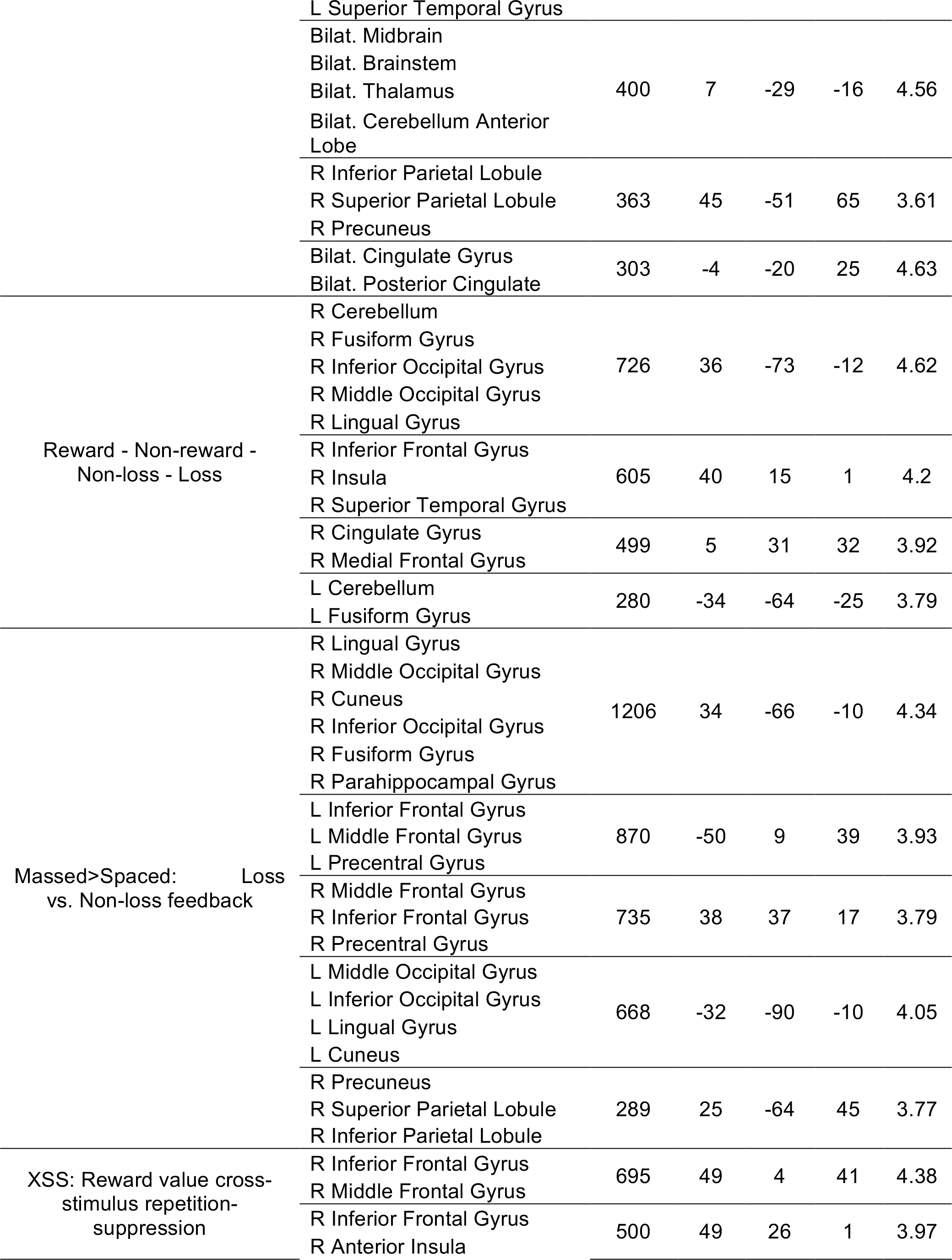
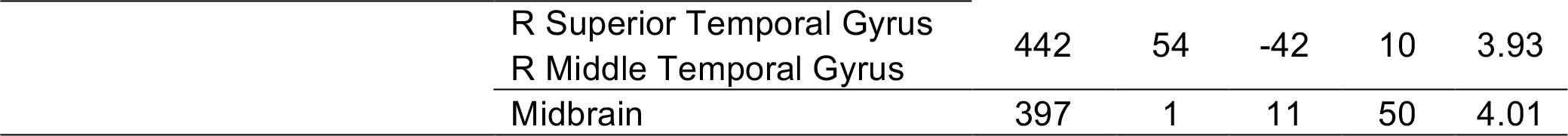
Summary of univariate analysis results. Clusters of activity exceeding whole-brain p < 0.05 FWE-corrected. Within each cluster, the first 10 regions are listed that include >= 10 voxels of a cluster.

To gain greater insight into the neural response to massed-and spaced-trained stimuli, we leveraged multivariate analysis methods. Specifically, we tested whether distributed patterns of brain activity within regions of interest or in a whole-brain searchlight analysis were able to discriminate between reward value, spaced vs. massed training condition, or their interaction. Our primary question was whether patterns of activity differentially discriminated the value of spaced-vs. massed-trained stimuli.

Our first analysis tested for patterns that discriminated between reward-vs. loss-associated stimuli. In the striatal region of interest, classification was not significantly different than zero (49.5 % CI [47.5 51.6]; t_(30)_ = −0.48, p = 0.63), and a similar null result was found in the hippocampus and parahippocampus MTL ROI (49.1 % CI [47.1 51.2]; t_(30)_ = −0.89, p = 0.38). Using a whole-brain searchlight analysis, thresholding at the standard cluster-forming threshold of p < 0.005 resulted in a large single cluster spanning much of the brain; for this reason, we used a more stringent cluster-forming threshold of p < 0.0005 in order to obtain more interpretable clusters. We identified several regions that showed significant value discrimination, including the left pre-and postcentral gyrus and a large bilateral cluster in the posterior and ventral occipital cortex (p < 0.05 whole-brain FWE-corrected; **Fig. 6**; Table 1).

**Figure 6.**
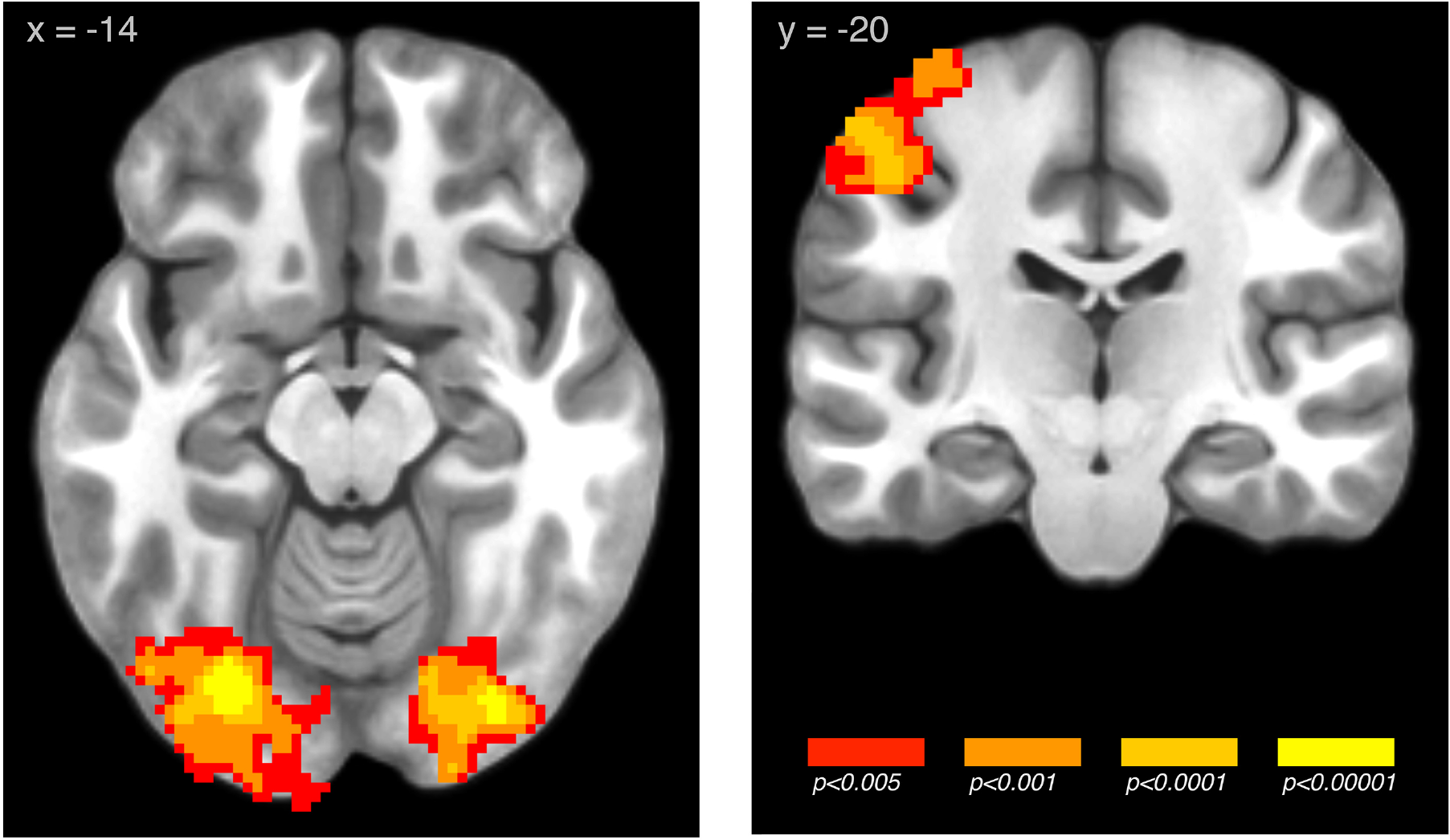
Searchlight pattern classification of reward-vs. loss-associated stimuli across the massed and spaced conditions (images whole-brain p < 0.05 FWE corrected; unthresholded map available at https://neurovault.org/images/59040/) For univariate results of the response to reward and loss feedback, see Figure 6-1.

**Figure 6-1.**
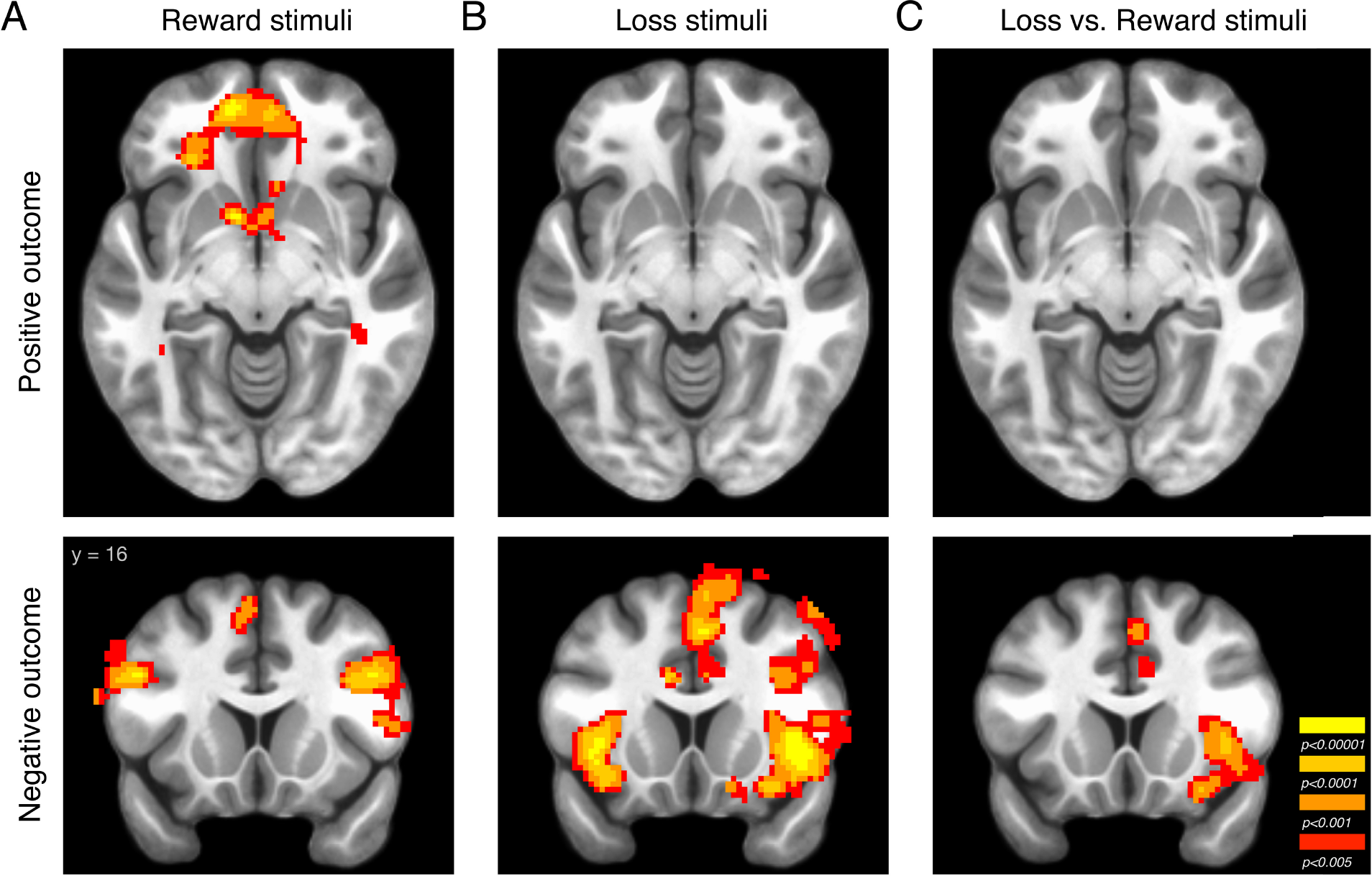
Univariate contrast of hit vs. miss feedback for reward-and loss-associated stimuli. ***A***, Reward-associated stimuli: activity related to hit > miss (top) and miss > hit (bottom). ***B***, Loss-associated stimuli: activity related to hit > miss (top) and miss > hit (bottom). ***C***, Contrast of loss-vs. reward-associated stimuli for hit > miss (top) and miss > hit (bottom).

To directly compare value-discriminating regions across condition, we examined the interaction of value by spacing condition. This analysis involved the contrast of two separate classifiers, one trained to discriminate reward-vs. loss-associated stimuli for massed-trained stimuli and the other for spaced-trained stimuli. In our ROI classification analysis, we found that patterns of activity in the MTL showed significantly stronger discrimination for spaced vs. massed values (difference, 8.2 % CI [3.8 12.5]; t_(30)_ = 3.81, p < 0.001; **Fig. 7A**). Importantly, the effect in the spaced condition alone was significant (55.5 % CI [53.1 57.9]; t_(30)_ = 4.58, p < 0.001; massed, 47.3 % CI [44.0 50.7]; t_(30)_ = - 1.60, p = 0.12). In the striatum, we found a similar effect (difference, 7.2 % CI [3.1 11.2]; t_(30)_ = 3.63, p = 0.001), but the difference is difficult to interpret given the below-chance performance in the massed condition (53.5 % CI [50.2 56.7]; t_(30)_ = 2.20, p = 0.036; massed, 46.3 % CI [43.8 48.8]; t_(30)_ = −3.01, p = 0.005).

Next, we examined whether more local patterns of activity showed significant discrimination of spaced-vs. massed-trained values. Thresholding at the standard cluster-forming threshold of p < 0.005 resulted in a large single cluster spanning much of the brain; as above, for this reason, we used a more stringent cluster-forming threshold of p < 0.0005 in order to obtain more interpretable clusters. We found multiple clusters exhibiting greater value discrimination in the spaced vs. massed condition, including the bilateral dorsolateral prefrontal cortex (DLPFC), the ventromedial prefrontal cortex (VMPFC), and orbitofrontal cortex (OFC) (**Fig. 7B**, Table 1). The searchlight analysis also demonstrated that the stronger classification of value observed in the spaced vs. massed conditions in the MTL ROI analysis were also found in the local searchlight analysis in the right hippocampus and parahippocampus (**Fig. 7C**; Table 1.). No regions showed greater discrimination of massed-trained values over spaced-trained values.

**Table 1.**
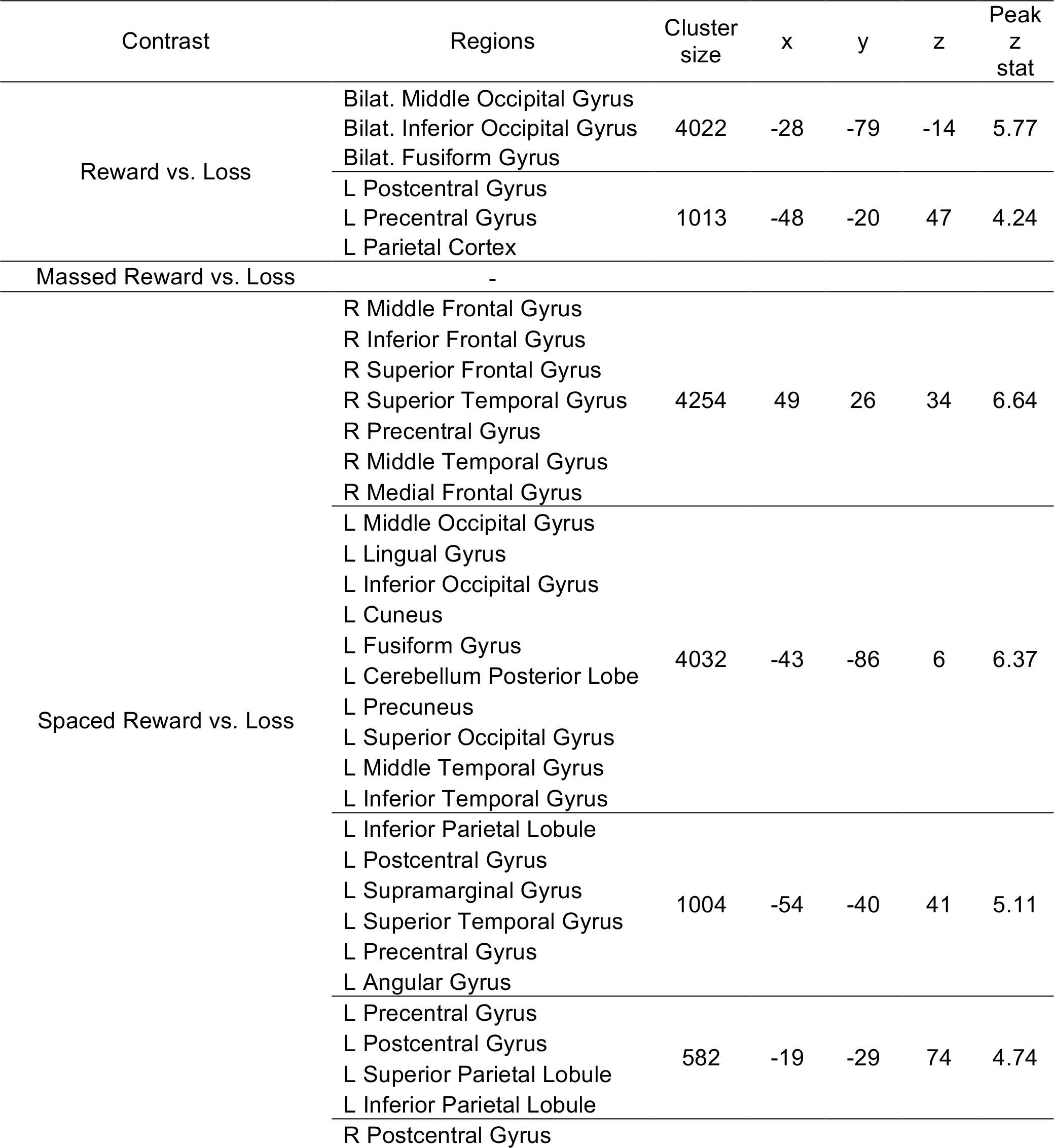
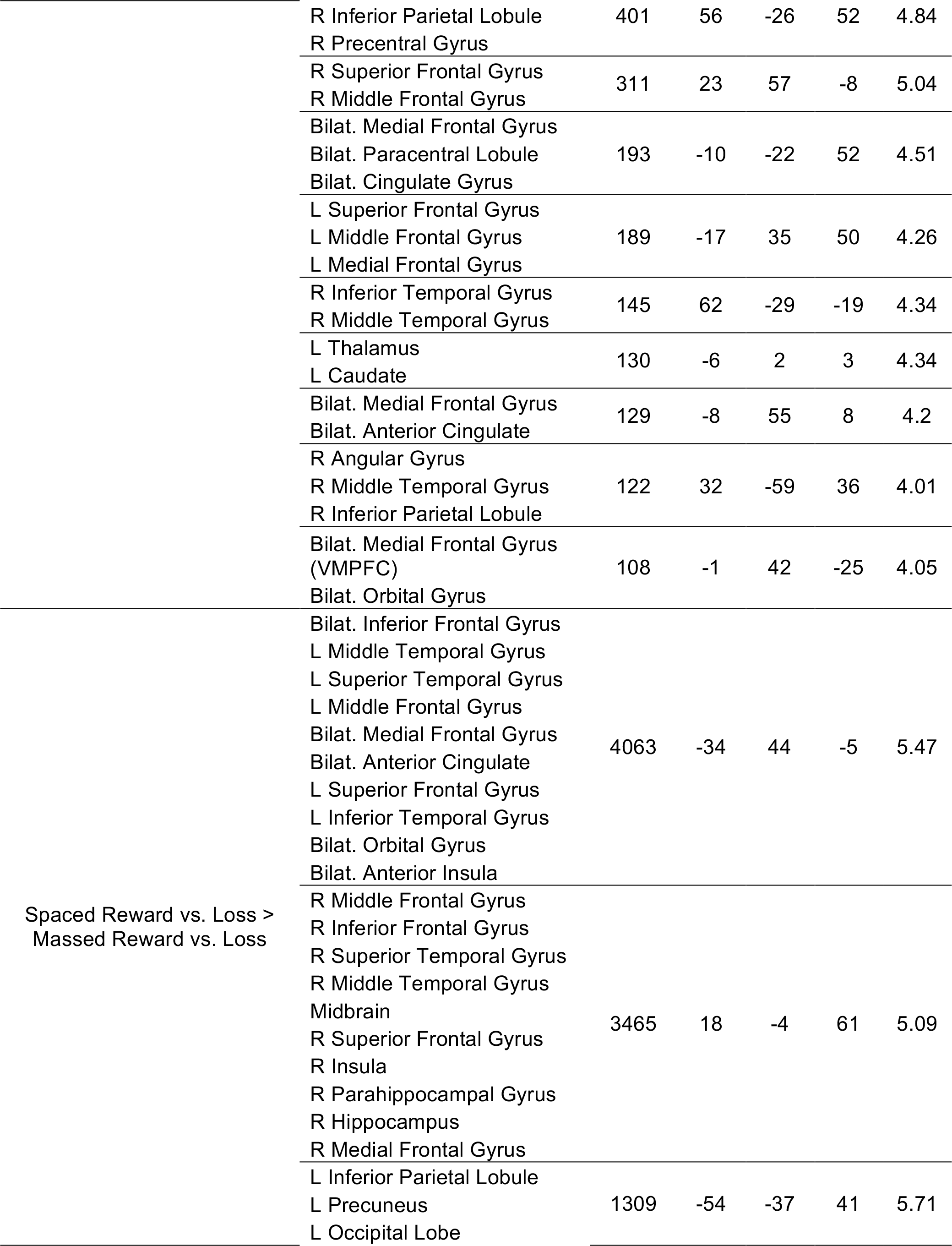
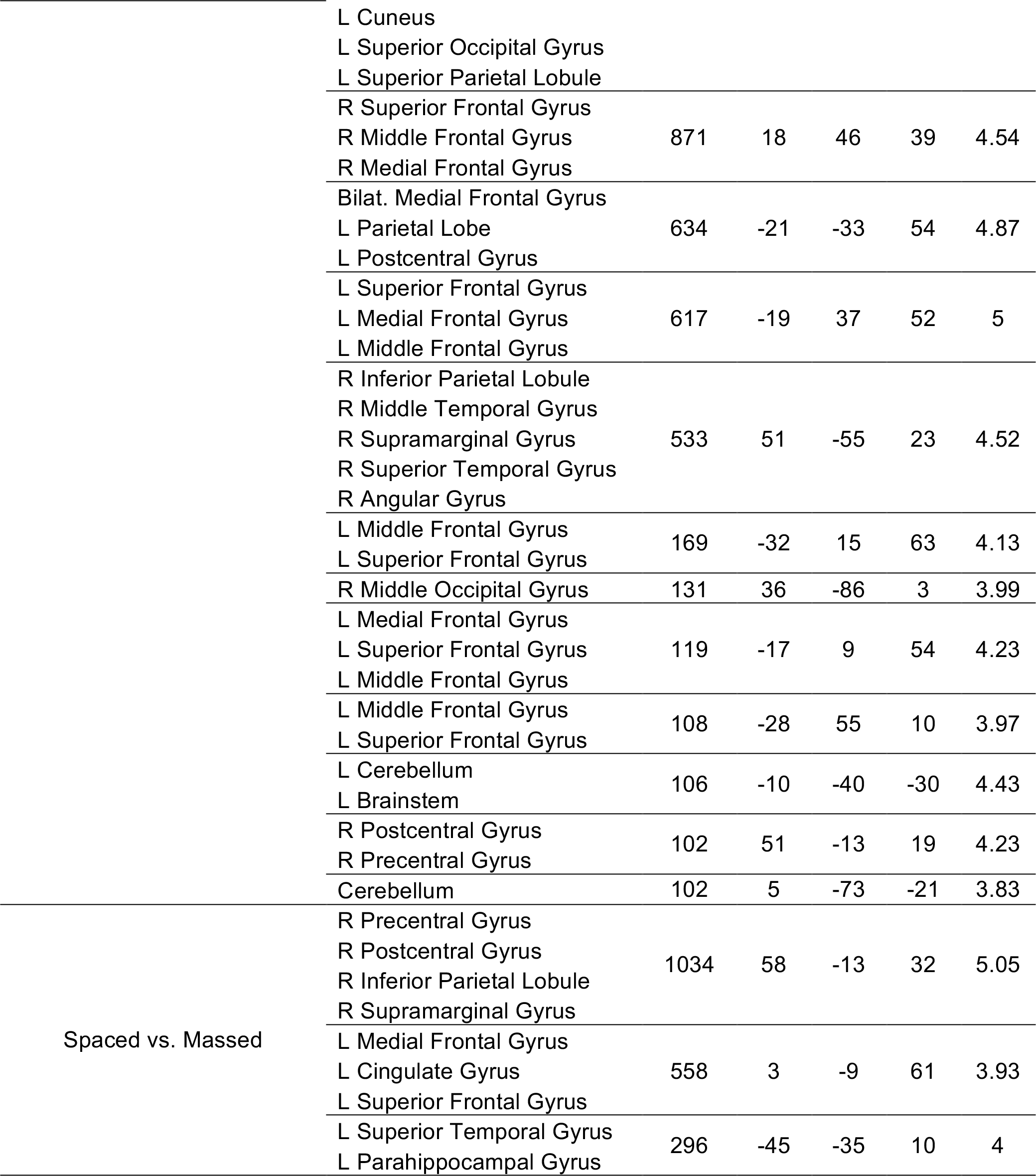
Summary of multivariate whole-brain searchlight analysis results. Clusters of activity exceeding whole-brain p < 0.05 FWE-corrected. Within each cluster, the first 10 regions are listed that include >= 10 voxels of a cluster. For the spaced value and spaced vs. massed value results, the cluster-forming threshold was increased to p < 0.005 to produce more interpretable clusters. For univariate results, see Table 1-1.

**Figure 7.**
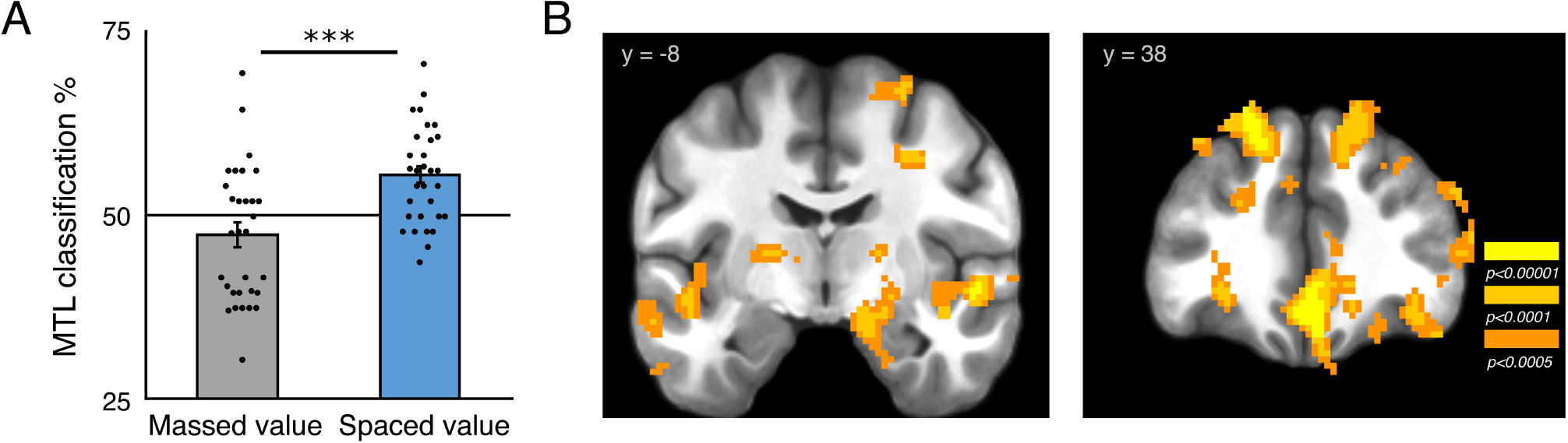
Pattern classification of spaced-trained values vs. massed-trained values. ***A***, MTL (hippocampus and parahippocampus) ROI shows significant classification of spaced versus massed values. ***B***, whole-brain searchlight analysis identified a cluster in the right medial temporal lobe. (*** p < 0.001; images whole-brain p < 0.05 FWE corrected; unthresholded map available at https://neurovault.org/images/59031/)

Finally, we examined the effect of spaced training by investigating which brain regions could successfully discriminate between spaced-vs. massed-trained stimuli. We found that the striatum showed significant discrimination of spacing condition (52.1 % CI [50.4 53.8]; t_(30)_ = 2.57, p = 0.016; **Fig. 8A**) while the effect in the MTL was not significant (51.1 % CI [49.5 52.6]; t_(30)_ = 1.40, p = 0.17). In the whole-brain searchlight analysis, we found several regions that discriminated the effect of time of training, including the left cingulate / supplementary motor area (3 −9 61; z = 3.93, p < 0.001 FWE; Table 1) and right pre-and post-central gyrus (58 −13 32; z = 5.05, p < 0.0001 FWE; **Fig. 8B**).

**Figure 8.**
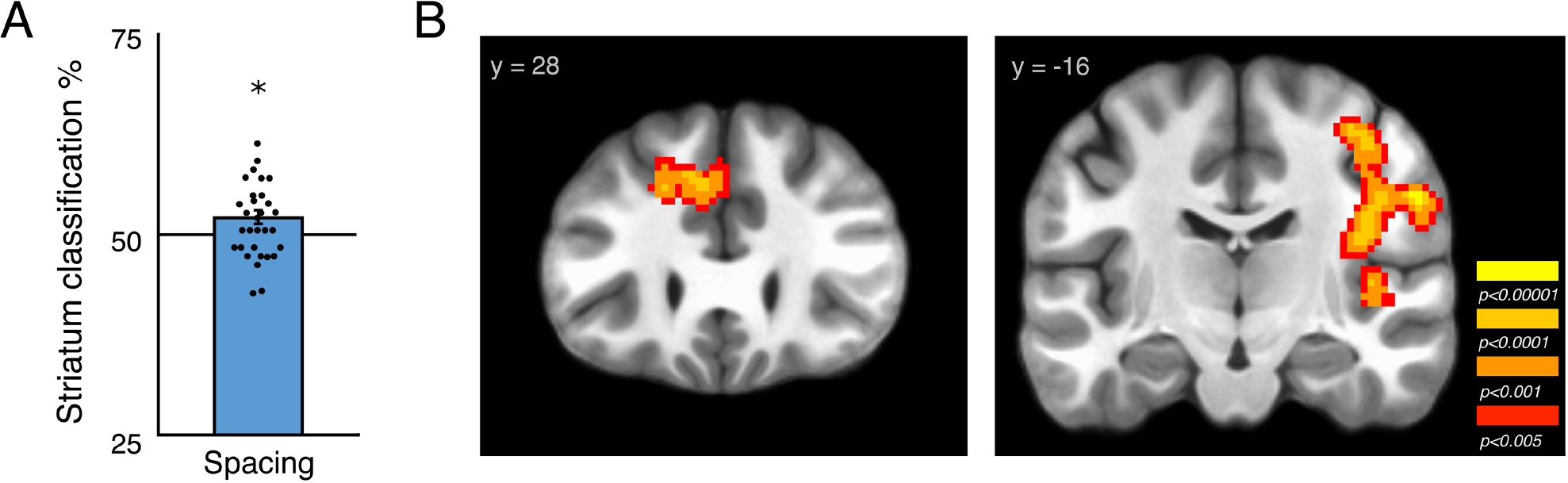
Pattern classification of spaced-vs. massed-trained stimuli. ***A***, Striatal ROI shows significant classification. ***B***, searchlight analysis identified additional clusters including the left cingulate and right pre-and post-central gyrus. (* p < 0.05; images whole-brain p < 0.05 FWE corrected; unthresholded map available at https://neurovault.org/images/59041/)

## Discussion

When reward-based learning is distributed over time instead of massed in a single session, we found significant gains in the long-term maintenance of learned value associations. Controlling for the amount of training as well as post-training performance, across two experiments we found that stimuli trained across weeks exhibited significantly stronger maintenance of value associations 3 weeks later. Conversely, single-session massed training, as commonly employed in human reward-based learning research, resulted in weaker maintenance of value associations. Decaying memory for massed-trained stimuli may be related to reliance on short-term memory to support massed learning, and supporting this view, we found that initial learning performance was significantly correlated with individual differences in working memory capacity.

Neurally, we found that distributed patterns of activity in the MTL and cortex discriminated between well-learned versus newly-learned value associations. Moreover, patterns of activity in the striatum discriminated well-learned versus newly-learned stimuli independent of value. These results were found in a task where participants learned the best of two responses for single reward-or loss-associated stimuli (similar to Pavlovian designs in non-human primates, e.g. Schultz et al., 1997; Kim and Hikosaka, 2013). It will be important for future research to verify that they extend to learning and maintenance of learned values in stable choice situations, such as a probabilistic selection task (Grogan et al., 2017). Together, these results indicate that reward associations acquired from weeks of training, in contrast to a single condensed session of learning, elicit stronger neural differentiation of value and may be more effective at guiding choices toward reward-associated options in the future.

Previous research has shown powerful effects of spacing in humans in memory and educational settings, following the initial work of Ebbinghaus (reported in Ebbinghaus, 1913; Cepeda et al., 2006). For reward-based learning, a beneficial effect of spacing has been well-established in other species (Teichner, 1952; Carew et al., 1972; Terrace et al., 1975). In humans, however, spacing has only been investigated in aversive eyeblink conditioning, which relies on a specialized cerebellar circuit (Humphreys, 1940; Spence and Norris, 1950; Kim and Thompson, 1997). Separately, effects of spacing have been investigated in category learning, which shares some structural similarities to the two-response instrumental task utilized in our experiments (Seger and Peterson, 2013; Carvalho and Goldstone, 2014). However, in contrast to the well-documented positive effects of spacing on feedback-based learning – where perceptual features do not allow generalization – category learning research indicates that the effect of alternating different examples benefits learning more than temporal spacing (Kang and Pashler, 2012).

From animal studies, reward-based learning is known to depend on the striatum and its midbrain dopaminergic projections (Schultz et al., 1997; Rangel et al., 2008; Steinberg et al., 2013). It is possible that condensed single-session learning in humans is primarily supported by the same neural mechanisms that support long-term learning. However, both our results and other recent findings strongly suggest that learning performance in tasks with condensed repetitions of stimuli additionally benefit from short-term cognitive mechanisms such as working memory (Collins and Frank, 2012; Collins et al., 2014). In human work on rapidly-paced paradigms, dopaminergic manipulations have been shown to affect performance (Frank et al., 2004; Pessiglione et al., 2006). While a full discussion of this research is beyond the scope of the present paper, we support the interpretation that some part of this shift is likely due to actions on a mechanism involving dopamine-induced synaptic plasticity in the striatum (though see Grogan et al., 2017). However, unlike work in animals where region and cell-type specific manipulations of dopaminergic and striatal neurons is possible (Steinberg et al., 2013; Ferenczi et al., 2016), pharmacological manipulations in humans have whole-brain effects. As dopamine also plays a significant role in higher cognitive functions including working memory, it is thus difficult to disentangle the effects of dopaminergic drugs on striatal plasticity and working memory processes (Cools, 2011; Matsumoto and Takada, 2013). A potential limitation of our experiments, as noted above, is that while our paradigm involves learning the value of single stimuli, similar to animal work on feedback-based learning (Schultz et al., 1997; Kim and Hikosaka, 2013; Ghazizadeh et al., 2018), it is not yet clear if these learning and maintenance results directly translate to a stable two-alternative bandit task.

Our results extend previous findings on the role of working memory in feedback learning by demonstrating that, in addition to a relationship with principal components of learning model fits (Collins et al., 2014), individual differences in working memory capacity are positively related to a simple measure of learning performance. The current experimental paradigm also includes both reward-and loss-associated stimuli, going beyond the conditional associative learning paradigm used previously (Petrides, 1985; Collins and Frank, 2012) and allowing for both forced-choice preference tests of learned values and an investigation into neural patterns that differentiate between learned values. Finally, our results indicate that more flexible models with multiple timescales of forgetting (and learning) may better account for the data than current models employing a single short-term working memory module (Collins and Frank, 2012).

What neural mechanisms support the improvement in long-term maintenance of values with spaced training? Our finding of significant classification of reward versus loss associations for spaced-trained but not massed-trained associations in the MTL, including the hippocampus, indicates a potentially novel role for the hippocampus in representing well-learned values. While the hippocampus is known to respond to reward and value (Lebreton et al., 2009; Wirth et al., 2009; Lee et al., 2012), hippocampal dysfunction does not eliminate the capacity of animals or humans to gradually learn the value of stimuli (e.g. Packard et al., 1989; Knowlton et al., 1996; Bayley et al., 2005). However, without the support of the hippocampus, feedback-based learning in humans is extraordinarily slow and inflexible (Bayley et al., 2005).

While often viewed as opposing systems, recent evidence suggests that striatal and hippocampal systems may cooperate during reward-based learning (Lansink et al., 2009; van der Meer et al., 2010; Foerde and Shohamy, 2011). Specifically, the MTL may support learning and decision making by acquiring statistical structure of stimulus-feedback associations (Schapiro et al., 2012) or by providing information about previous episodes (Shadlen and Shohamy, 2016), a proposal supported by recent research (Murty et al., 2016; Wimmer and Buechel, 2016; Bornstein et al., 2017). The MTL may play a larger role in supporting learning over longer timescales, allowing for learning across contexts as well as the consolidation of synaptic plasticity (Kramar et al., 2012; Aziz et al., 2014; Smolen et al., 2016), which could explain selective value discrimination in the hippocampus for spaced-but not massed-trained associations. Computationally, spaced training may allow for the benefits of offline replay as employed in models such as DYNA, where model-free values are trained by post-event replay of experience (Sutton, 1990; Johnson and Redish, 2005; Gershman et al., 2014; Russek et al., 2017).

Additionally, we found that patterns of activity in the striatum discriminated spaced-trained versus massed-trained stimuli overall. Decades of animal research have shown that different regions of the striatum are important for different types of reward associations, with the dorsomedial striatum critical for flexible (and newly-acquired) goal-directed learning and the dorsolateral striatum critical for inflexible model-free and habit learning (Balleine and Dickinson, 1998; Yin and Knowlton, 2006; Kim and Hikosaka, 2013; Foerde, 2018). In contrast to previous fMRI studies that employed a multi-day design (Tricomi et al., 2009; Wunderlich et al., 2012), our experimental design allows for a direct comparison between equivalent amounts of spaced and massed training. We did not find any effect of spacing on univariate measures of value in the striatum, in contrast to previous studies (Tricomi et al., 2009; Wunderlich et al., 2012), although null results should be treated with caution. Recent findings in non-human primates indicate that a novel population of striatum-projecting dopamine neurons responds to well-learned value associations, even after stimulus-reward associations are extinguished (Kim et al., 2015). Such a neural mechanism may support a “habit” of attentional orientation to reward-associated that is resistant to extinction (Kim et al., 2015; Anderson, 2016). We did not collect a measure of devaluation sensitivity, the classic test of habitual behavior - albeit one difficult to administer in humans (Dickinson, 1985; Graybiel, 2008; Tricomi et al., 2009; de Wit et al., in press). Whether or not the learned stimulus-action associations remained sensitive to outcomes, our results indicate that the brain may retain the ability to remember and recall the value associated with stimulus using other representations in memory, such as those supported by the MTL.

Our results have implications for understanding reward-based learning in the healthy brain and for translating this research to patient populations (Huys et al., 2016) and more ecologically valid experimental designs (Moutoussis et al., 2016). The interpretation of parameters derived from massed feedback learning paradigms is difficult for various reasons, including, as we demonstrate, the contribution of working memory to performance (see also Collins and Frank, 2012). Additionally, the decaying nature of value associations learned in massed-training tasks suggests that parameters
derived from massed paradigms may not translate to how people acquire lasting value associations and habits over time outside the lab. Our experiments suggest that the long-term maintenance of value associations may be a promising individual difference measure to explore in future studies. Finally, our experimental design provides a starting point for testing how over-learned value associations may be unlearned, with implications for research on behavioral change.

In summary, across two studies we found that spacing of condensed sessions of reward-based learning across weeks resulted in significantly greater maintenance of conditioned value associations than training across minutes. Our experiments represent the first demonstration of spacing effects on reward-based learning in humans and identify neural signatures specific to well-learned vs. transient value associations in the human brain. Overall, our results indicate that spaced reward-based learning and long-term maintenance of conditioning may provide cleaner measures of feedback-based learning than current measures. This possibility has implications for the interpretation and direction of reward-based learning research, as feedback learning paradigms are becoming widely used in studies of mood and psychiatric disorders as well as addiction (Herbener, 2009; Maia and Frank, 2011; Montague et al., 2012; Whitton et al., 2015).

## Acknowledgments

The authors thank Patrick Bissett, Ross Blair, and Oscar Esteban. Research was supported by NIH R01AG041653 [RP], a research fellowship from the Deutsche Forschungsgemeinschaft [GEW] and a pilot seed grant from the Stanford Center for Cognitive and Neurobiological Imaging [GEW].

## References

Anderson BA (2016) The attention habit: how reward learning shapes attentional selection. Ann N Y Acad Sci 1369:24–39.

Andersson JL, Skare S, Ashburner J (2003) How to correct susceptibility distortions in spin-echo echo-planar images: application to diffusion tensor imaging. Neuroimage 20:870–888.

Avants BB, Epstein CL, Grossman M, Gee JC (2008) Symmetric diffeomorphic image registration with cross-correlation: evaluating automated labeling of elderly and neurodegenerative brain. Med Image Anal 12:26–41.

Aziz W, Wang W, Kesaf S, Mohamed AA, Fukazawa Y, Shigemoto R (2014) Distinct kinetics of synaptic structural plasticity, memory formation, and memory decay in massed and spaced learning. Proc Natl Acad Sci U S A 111:E194–202.

Balleine BW, Dickinson A (1998) Goal-directed instrumental action: contingency and incentive learning and their cortical substrates. Neuropharmacology 37:407–419.

Barron HC, Dolan RJ, Behrens TE (2013) Online evaluation of novel choices by simultaneous representation of multiple memories. Nat Neurosci 16:1492–1498.

Barron HC, Garvert MM, Behrens TE (2016) Repetition suppression: a means to index neural representations using BOLD? Philos Trans R Soc Lond B Biol Sci 371.

Bartra O, McGuire JT, Kable JW (2013) The valuation system: a coordinate-based meta-analysis of BOLD fMRI experiments examining neural correlates of subjective value. Neuroimage 76:412–427.

Bayley PJ, Frascino JC, Squire LR (2005) Robust habit learning in the absence of awareness and independent of the medial temporal lobe. Nature 436:550–553.

Bornstein AM, Khaw MW, Shohamy D, Daw ND (2017) Reminders of past choices bias decisions for reward in humans. Nat Commun 8:15958.

Brainard DH (1997) The Psychophysics Toolbox. Spat Vis 10:433–436.

Carew TJ, Pinsker HM, Kandel ER (1972) Long-term habituation of a defensive withdrawal reflex in aplysia. Science 175:451–454.

Carvalho PF, Goldstone RL (2014) Effects of interleaved and blocked study on delayed test of category learning generalization. Frontiers in psychology 5.

Cepeda NJ, Pashler H, Vul E, Wixted JT, Rohrer D (2006) Distributed practice in verbal recall tasks: A review and quantitative synthesis. Psychol Bull 132:354–380.

Chang CC, Lin C (2011) LIBSVM: a library for support vector machines. ACM Trans Intell Syst Technol 2:1–27.

Colas JT, Pauli WM, Larsen T, Tyszka JM, O’Doherty JP (2017) Distinct prediction errors in mesostriatal circuits of the human brain mediate learning about the values of both states and actions: evidence from high-resolution fMRI. PLoS Comput Biol 13:e1005810.

Cole MW, Laurent P, Stocco A (2013) Rapid instructed task learning: a new window into the human brain’s unique capacity for flexible cognitive control. Cogn Affect Behav Neurosci 13:1–22.

Collins AG, Frank MJ (2012) How much of reinforcement learning is working memory, not reinforcement learning? A behavioral, computational, and neurogenetic analysis. Eur J Neurosci 35:1024–1035.

Collins AG, Brown JK, Gold JM, Waltz JA, Frank MJ (2014) Working memory contributions to reinforcement learning impairments in schizophrenia. J Neurosci 34:13747–13756.

Cools R (2011) Dopaminergic control of the striatum for high-level cognition. Curr Opin Neurobiol 21:402–407.

Cox RW (1996) AFNI: software for analysis and visualization of functional magnetic resonance neuroimages. Comput Biomed Res 29:162–173.

Dale AM, Fischl B, Sereno MI (1999) Cortical surface-based analysis. I. Segmentation and surface reconstruction. Neuroimage 9:179–194.

Daw ND, O’Doherty JP, Dayan P, Seymour B, Dolan RJ (2006) Cortical substrates for exploratory decisions in humans. Nature 441:876–879.

de Leeuw JR (2015) jsPsych: a JavaScript library for creating behavioral experiments in a Web browser. Behav Res Methods 47:1–12.

de Wit S, Kindt M, Knot SL, Verhoeven AAC, Robbins TW, Gasull-Camos J, Evans M, Mirza H, Gillan CM (in press) Shifting the balance between goals and habits: five failures in experimental habit induction. J Exp Psychol Gen.

Deichmann R, Gottfried JA, Hutton C, Turner R (2003) Optimized EPI for fMRI studies of the orbitofrontal cortex. Neuroimage 19:430–441.

Dickerson KC, Li J, Delgado MR (2011) Parallel contributions of distinct human memory systems during probabilistic learning. Neuroimage 55:266–276.

Dickinson A (1985) Actions and habits: the development of behavioural autonomy. Phil Trans R Soc Lond B 308:67–78.

Dolan RJ, Dayan P (2013) Goals and habits in the brain. Neuron 80:312–325.

Ebbinghaus H (1913) Memory: A contribution to experimental psychology.

Eichenbaum H, Cohen NJ (2001) From Conditioning to Conscious Recollection: Memory Systems of the Brain. New York: Oxford University Press.

Eldar E, Roth C, Dayan P, Dolan RJ (2018) Decodability of Reward Learning Signals Predicts Mood Fluctuations. Current biology : CB.

Ferenczi EA, Zalocusky KA, Liston C, Grosenick L, Warden MR, Amatya D, Katovich K, Mehta H, Patenaude B, Ramakrishnan C, Kalanithi P, Etkin A, Knutson B, Glover GH, Deisseroth K (2016) Prefrontal cortical regulation of brainwide circuit dynamics and reward-related behavior. Science 351:aac9698.

Fiorillo CD, Newsome WT, Schultz W (2008) The temporal precision of reward prediction in dopamine neurons. Nat Neurosci 11:966–973.

Foerde K (2018) What are habits and do they depend on the striatum? A view from the study of neuropsychological populations. Curr Opin Beh Sci 20:17–24.

Foerde K, Shohamy D (2011) Feedback timing modulates brain systems for learning in humans. J Neurosci 31:13157–13167.

Foerde K, Race E, Verfaellie M, Shohamy D (2013) A role for the medial temporal lobe in feedback-driven learning: evidence from amnesia. J Neurosci 33:5698–5704.

Fonov VS, Evans AC, McKinstry RC, Almli CR, Collins DL (2009) Unbiased Nonlinear Average Age-Appropriate Brain Templates from Birth to Adulthood. Neuroimage 47.

Frank MJ, Seeberger LC, O’Reilly R C (2004) By carrot or by stick: cognitive reinforcement learning in parkinsonism. Science 306:1940–1943.

Friston KJ, Worsley KJ, Frackowiak SJ, Mazziotta JC, Evans AC (1993) Assessing the significance of focal activations using their spatial extent. Human Brain Mapping 1:210–220.

Gerraty RT, Davidow JY, Wimmer GE, Kahn I, Shohamy D (2014) Transfer of learning relates to intrinsic connectivity between hippocampus, ventromedial prefrontal cortex, and large-scale networks. J Neurosci 34:11297–11303.

Gershman SJ, Markman AB, Otto AR (2014) Retrospective revaluation in sequential decision making: a tale of two systems. J Exp Psychol Gen 143:182–194.

Ghazizadeh A, Griggs W, Leopold DA, Hikosaka O (2018) Temporal-prefrontal cortical network for discrimination of valuable objects in long-term memory. Proc Natl Acad Sci U S A 115:E2135–E2144.

Gorgolewski K, Burns CD, Madison C, Clark D, Halchenko YO, Waskom ML, Ghosh SS (2011) Nipype: a flexible, lightweight and extensible neuroimaging data processing framework in python. Frontiers in neuroinformatics 5:13.

Graybiel AM (2008) Habits, rituals, and the evaluative brain. Annu Rev Neurosci 31:359–387.

Greve DN, Fischl B (2009) Accurate and robust brain image alignment using boundary-based registration. Neuroimage 48:63–72.

Grogan JP, Tsivos D, Smith L, Knight BE, Bogacz R, Whone A, Coulthard EJ (2017) Effects of dopamine on reinforcement learning and consolidation in Parkinson’s disease. Elife 6.

Hebart MN, Gorgen K, Haynes JD (2014) The Decoding Toolbox (TDT): a versatile software package for multivariate analyses of functional imaging data. Frontiers in neuroinformatics 8:88.

Herbener ES (2009) Impairment in long-term retention of preference conditioning in schizophrenia. Biol Psychiatry 65:1086–1090.

Houk JC, Adams JL, Barto AG (1995) A model of how the basal ganglia generate and use neural signals that predict reinforcement. In: Models of information processing in the basal ganglia (Houk JC, Davis JL, Beiser DG, eds), pp 249–270. Cambridge, MA: MIT Press.

Humphreys LG (1940) Distributed practice in the development of the conditioned eyelid response. J Gen Psychol 22:379–385.

Huys QJ, Maia TV, Frank MJ (2016) Computational psychiatry as a bridge from neuroscience to clinical applications. Nat Neurosci 19:404–413.

Jenkinson M (2003) Fast, automated, N-dimensional phase-unwrapping algorithm. Magn Reson Med 49:193–197.

Jenkinson M, Bannister P, Brady M, Smith S (2002) Improved optimization for the robust and accurate linear registration and motion correction of brain images. Neuroimage 17:825–841.

Johnson A, Redish AD (2005) Hippocampal replay contributes to within session learning in a temporal difference reinforcement learning model. Neural Netw 18:1163–1171.

Kang SHK, Pashler H (2012) Learning Painting Styles: Spacing is Advantageous when it Promotes Discriminative Contrast. Appl Cognitive Psych 26:97–103.

Kim HF, Hikosaka O (2013) Distinct basal ganglia circuits controlling behaviors guided by flexible and stable values. Neuron 79:1001–1010.

Kim HF, Ghazizadeh A, Hikosaka O (2015) Dopamine Neurons Encoding Long-Term Memory of Object Value for Habitual Behavior. Cell 163:1165–1175.

Kim JJ, Thompson RF (1997) Cerebellar circuits and synaptic mechanisms involved in classical eyeblink conditioning. Trends Neurosci 20:177–181.

Klein-Flugge MC, Barron HC, Brodersen KH, Dolan RJ, Behrens TE (2013) Segregated encoding of reward-identity and stimulus-reward associations in human orbitofrontal cortex. J Neurosci 33:3202–3211.

Knowlton BJ, Mangels JA, Squire LR (1996) A neostriatal habit learning system in humans. Science 273:1399–1402.

Kramar EA, Babayan AH, Gavin CF, Cox CD, Jafari M, Gall CM, Rumbaugh G, Lynch G (2012) Synaptic evidence for the efficacy of spaced learning. Proc Natl Acad Sci U S A 109:5121–5126.

Lakens D (2017) Equivalence tests: A practical primer for t-tests, correlations, and metaanalyses. Soc Psychol Personal Sci.

Lansink CS, Goltstein PM, Lankelma JV, McNaughton BL, Pennartz CM (2009) Hippocampus leads ventral striatum in replay of place-reward information. PLoS Biol 7:e1000173.

Lebreton M, Jorge S, Michel V, Thirion B, Pessiglione M (2009) An automatic valuation system in the human brain: evidence from functional neuroimaging. Neuron 64:431–439.

Lee H, Ghim JW, Kim H, Lee D, Jung M (2012) Hippocampal neural correlates for values of experienced events. J Neurosci 32:15053–15065.

Lewandowsky S, Oberauer K, Yang LX, Ecker UK (2010) A working memory test battery for MATLAB. Behav Res Methods 42:571–585.

Maia TV, Frank MJ (2011) From reinforcement learning models to psychiatric and neurological disorders. Nat Neurosci 14:154–162.

Matsumoto M, Takada M (2013) Distinct representations of cognitive and motivational signals in midbrain dopamine neurons. Neuron 79:1011–1024.

Montague PR, Dolan RJ, Friston KJ, Dayan P (2012) Computational psychiatry. Trends Cogn Sci 16:72–80.

Moutoussis M, Eldar E, Dolan RJ (2016) Building a New Field of Computational Psychiatry. Biol Psychiatry.

Murty VP, FeldmanHall O, Hunter LE, Phelps EA, Davachi L (2016) Episodic memories predict adaptive value-based decision-making. J Exp Psychol Gen 145:548–558.

O’Doherty JP, Dayan P, Friston K, Critchley H, Dolan RJ (2003) Temporal difference models and reward-related learning in the human brain. Neuron 38:329–337.

Otto AR, Raio CM, Chiang A, Phelps EA, Daw ND (2013) Working-memory capacity protects model-based learning from stress. Proc Natl Acad Sci U S A 110:20941–20946.

Packard MG, Hirsh R, White NM (1989) Differential effects of fornix and caudate nucleus lesions on two radial maze tasks: evidence for multiple memory systems. J Neurosci 9:1465–1472.

Pessiglione M, Seymour B, Flandin G, Dolan RJ, Frith CD (2006) Dopamine-dependent prediction errors underpin reward-seeking behaviour in humans. Nature 442:1042–1045.

Petrides M (1985) Deficits on conditional associative-learning tasks after frontal-and temporal-lobe lesions in man. Neuropsychologia 23:601–614.

Plassmann H, O’Doherty J, Rangel A (2007) Orbitofrontal cortex encodes willingness to pay in everyday economic transactions. J Neurosci 27:9984–9988.

Power JD, Mitra A, Laumann TO, Snyder AZ, Schlaggar BL, Petersen SE (2014) Methods to detect, characterize, and remove motion artifact in resting state fMRI. Neuroimage 84:320–341.

Rangel A, Camerer C, Montague PR (2008) A framework for studying the neurobiology of value-based decision making. Nat Rev Neurosci 9:545–556.

Russek EM, Momennejad I, Botvinick MM, Gershman SJ, Daw ND (2017) Predictive representations can link model-based reinforcement learning to model-free mechanisms. PLoS Comput Biol 13:e1005768.

Schapiro AC, Kustner LV, Turk-Browne NB (2012) Shaping of object representations in the human medial temporal lobe based on temporal regularities. Current biology : CB 22:1622–1627.

Schapiro AC, Gregory E, Landau B, McCloskey M, Turk-Browne NB (2014) The necessity of the medial temporal lobe for statistical learning. J Cogn Neurosci 26:1736–1747.

Schuirmann DJ (1987) A comparison of the two one-sided tests procedure and the power approach for assessing the equivalence of average bioavailability. J Pharmacokinet Biopharm 15:657–680.

Schultz W (2011) Potential vulnerabilities of neuronal reward, risk, and decision mechanisms to addictive drugs. Neuron 69:603–617.

Schultz W, Dayan P, Montague PR (1997) A neural substrate of prediction and reward. Science 275:1593–1599.

Seger CA, Peterson EJ (2013) Categorization = decision making + generalization. Neuroscience and biobehavioral reviews 37:1187–1200.

Shadlen MN, Shohamy D (2016) Decision Making and Sequential Sampling from Memory. Neuron 90:927–939.

Smolen P, Zhang Y, Byrne JH (2016) The right time to learn: mechanisms and optimization of spaced learning. Nat Rev Neurosci 17:77–88.

Sochat VV, Eisenberg IW, Enkavi AZ, Li J, Bissett PG, Poldrack RA (2016) The Experiment Factory: Standardizing Behavioral Experiments. Frontiers in psychology 7:610.

Spence KW, Norris EB (1950) Eyelid conditioning as a function of the inter trial interval. Journal of Experimental Psychology 40:716–720.

Steinberg EE, Keiflin R, Boivin JR, Witten IB, Deisseroth K, Janak PH (2013) A causal link between prediction errors, dopamine neurons and learning. Nat Neurosci 16:966–973.

Sutton RS (1990) Integrated architectures for learning, planning, and reacting based on approximating dynamic programming. In: Proceedings of the Seventh International Conference on Machine Learning (Porter BW, Mooney RJ, eds), pp 216–224: Morgan Kaufmann.

Tanaka SC, Doya K, Okada G, Ueda K, Okamoto Y, Yamawaki S (2004) Prediction of immediate and future rewards differentially recruits cortico-basal ganglia loops. Nat Neurosci 7:887–893.

Teichner WH (1952) Experimental extinction as a function of the intertrial intervals during conditioning and extinction. Journal of Experimental Psychology 44:170–178.

Terrace HS, Gibbon J, Farrell L, Baldock MD (1975) Temporal factors influencing acquisition and maintenance of an autoshaped keypeck. Animal Learning & Behavior 3:53–62.

Tricomi E, Balleine BW, O’Doherty JP (2009) A specific role for posterior dorsolateral striatum in human habit learning. Eur J Neurosci 29:2225–2232.

Tustison NJ, Avants BB, Cook PA, Zheng Y, Egan A, Yushkevich PA, Gee JC (2010) N4ITK: improved N3 bias correction. IEEE Trans Med Imaging 29:1310–1320.

Tzourio-Mazoyer N, Landeau B, Papathanassiou D, Crivello F, Etard O, Delcroix N, Mazoyer B, Joliot M (2002) Automated anatomical labeling of activations in SPM using a macroscopic anatomical parcellation of the MNI MRI single-subject brain. Neuroimage 15:273–289.

van der Meer MA, Johnson A, Schmitzer-Torbert NC, Redish AD (2010) Triple Dissociation of Information Processing in Dorsal Striatum, Ventral Striatum, and Hippocampus on a Learned Spatial Decision Task. Neuron 67:25–32.

Whitton AE, Treadway MT, Pizzagalli DA (2015) Reward processing dysfunction in major depression, bipolar disorder and schizophrenia. Curr Opin Psychiatry 28:7–12.

Wimmer GE, Shohamy D (2012) Preference by association: how memory mechanisms in the hippocampus bias decisions. Science 338:270–273.

Wimmer GE, Buechel C (2016) Reactivation of reward-related patterns from single past episodes supports memory-based decision making. J Neurosci 36:2868–2880.

Wimmer GE, Daw ND, Shohamy D (2012) Generalization of value in reinforcement learning by humans. Eur J Neurosci 35:1092–1104.

Wimmer GE, Braun EK, Daw ND, Shohamy D (2014) Episodic memory encoding interferes with reward learning and decreases striatal prediction errors. J Neurosci 34:14901–14912.

Wirth S, Avsar E, Chiu CC, Sharma V, Smith AC, Brown E, Suzuki WA (2009) Trial outcome and associative learning signals in the monkey hippocampus. Neuron 61:930–940.

Wunderlich K, Dayan P, Dolan RJ (2012) Mapping value based planning and extensively trained choice in the human brain. Nat Neurosci 15:786–791.

Yin HH, Knowlton BJ (2006) The role of the basal ganglia in habit formation. Nat Rev Neurosci 7:464–476.

